# Modulating radical propagation in proteins by proton-coupled electron transfer and hydrogen bonding

**DOI:** 10.64898/2026.03.14.711208

**Authors:** Rebecca K. Zawistowski, Timothée Chauviré, Sutanuka Manna, Nandini Ananth, Brian R. Crane

## Abstract

Long-range protein electron transfer (ET) often depends on tryptophan and tyrosine residues acting as radical relay sites. For example, cytochrome c peroxidase (CcP) generates a W191^•+^ radical to increase ET from cytochrome c (Cc) to the active center. W191 substitution to Tyr reduces ET rates, but introduction of an adjacent general base (as Glu or His) at position 232 (Y191:E/H232 CcP) recovers activity. E232 fluorination lowers the p*K*a of the conjugate base and confirms that a hydrogen bond is critical to elevate the Y191^•^ formal potential for effective ET. Photoinitiated ET between Zn-porphyrin (ZnP) CcP (ZnCcP) and Cc also depends on activating Y191 with a basic residue, but through a different mechanism than for the peroxide-driven system. In ZnCcP, pH dependencies and solvent isotope effects indicate that proton-coupled electron transfer to the basic residue and ZnP^•+^, respectively, facilitates Y191^•^ formation. Replacing Cc with the irreversible oxidant [Co(NH3)5Cl]^2+^ isolates distinct protein radicals for characterization by Electron Paramagnetic Resonance (EPR) spectroscopy. Radical distributions and computation indicate that W191^•+^ lies close in potential to ZnP^•+^ and that the two radicals exchange on a slow time scale despite their close separation. Remarkably, Y191:E/H232 ZnCcP variants propagate radicals differently to peripheral sites depending on the nature of the 232 residue. QM/MM calculations support radical exchange between ZnP^•+^/Trp^•+^ and the importance of a hydrogen bond to Y191^•^ for maintaining a high potential to oxidize peripheral donors. These resolved reactivity patterns of CcP/ZnCcP have general relevance for engineering proton management to separate and migrate charge in proteins and potentially other molecular systems.

**For Table of Contents Only:** 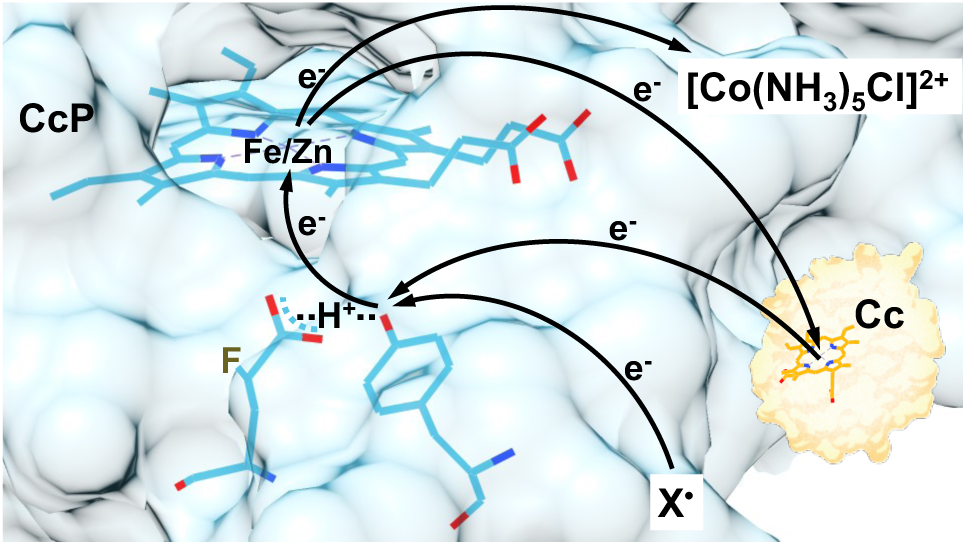

## Introduction

Protein long-range electron transfer (ET) is a centrally important process to many metabolic pathways including cellular respiration and photosynthesis [1–3]. In these long-range reactions, electrons transfer through protein matrices via the oxidation and reduction of redox active cofactors often separated by >10 Å or more. Despite the long separations, these ET reactions can still occur at remarkably fast rates, exceptional specificity, and with little loss in free energy [1–4]. Redox active protein residues, primarily tryptophan and tyrosine, facilitate multi-step ET by acting as oxidizable hole-hopping relay sites [4,5]. For example, Trp and Tyr radical intermediates migrate charge over distances that can span over 30 Å in systems such as ribonucleotide reductases (RNR), cryptochromes, BLUF photoreceptors, peroxidases, and photosystem II (PSII) [6–9]. Despite the ability of high potential metallocenters or photoexcited cofactors to oxidize either Trp or Tyr, these residues are often not interchangeable owing to their different sensitivities to local environment and the different p*K*as of their radical cations. Indeed, Proton-Coupled ET (PCET) plays a critical, complex and still understudied role in hole-hopping through Trp and Tyr [10,11]. For example, in PSII, PCET reactions of TyrZ and a nearby His residue allow P680^+^ to efficiently transfer electron holes to the oxygen evolving complex for water splitting [12–14]. Glu52 in the ribonucleotide reductase β-subunit has been shown to be essential for mediating charge propagation by Tyr radicals; however, several interpretations have been given for the underlying mechanism [15,16]. Furthermore, multistep ET reactions involving Trp/Tyr also have the potential to generate spin-correlated radical pairs (SCRPs) with relatively long phase-memory times [17]. Such SCRPs have the potential to be exquisitely sensitive detectors of local environment and have been implicated in, for example, the avian magnetic compass [18]. However, Trp and Tyr radicals have high redox potentials, and proteins provide many reactive sites. Thus, controlling reactivity is non-trivial. Investigating how local environment modulates relay residue redox chemistry will help us better understand mechanisms that underlie fundamental biological sensing and energy conversion.

**Scheme 1.**
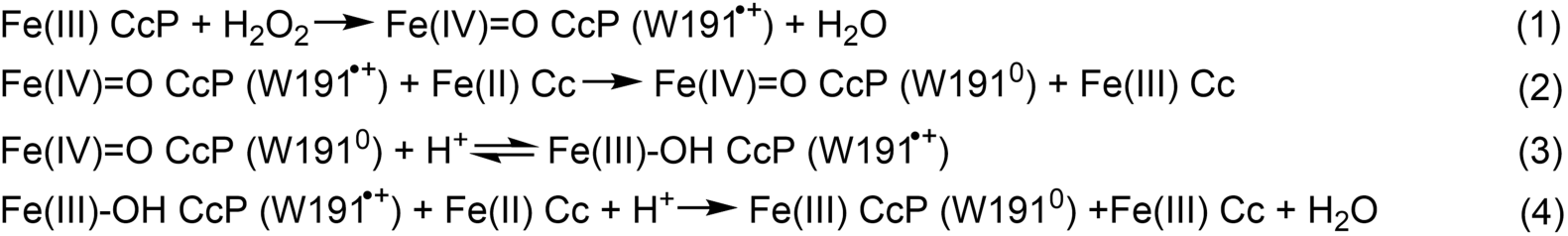
Reaction mechanism for Fe(III) CcP and Fe(II) Cc with H2O2

The complex formed between yeast cytochrome c peroxidase (CcP) and cytochrome c (Cc) has long stood as a model system for understanding long-range interprotein ET [19–23]. Moreover, it contains one of the most thoroughly characterized Trp radicals, which has been analyzed in detail by EPR spectroscopy [22,24–26]. The heme-containing CcP (FeCcP) reacts with H2O2 to form Compound I (Cpd I), which contains the ferryl heme and a second oxidized equivalent on Trp191 (W191, Scheme 1). Fe(IV)=O CcP is then reduced sequentially by two equivalents of Fe(II) Cc. W191^•+^ in Cpd I oxidizes the Fe(II) Cc heme which is 22 Å away (**Figure 1a**). Substitution of W191 to Gly or Phe (G/F191 CcP) eliminates CcP activity. Surprisingly, switching W191 for another oxidizable residue, Tyr191 (which we designate as the “Y191 CcP” variant) also renders CcP inactive, despite Y191 also forming a Cpd I-like state with peroxide [27]. Nonetheless, the activity of Y191 CcP can be partially rescued by introduction of an adjacent conjugate base through substitution of residue Leu232 in CcP for Glu or His (which we designate as Y191:E232 CcP or Y191:H232 CcP, respectively) [28]. These residues are both able to accept a proton from Tyr upon oxidation and facilitate hydrogen bonding to the formed radical [12–14,29]. Crystal structures of the Y191, Y191:E232 and Y191:H232 variants reveal that the E/H232 residues are in close proximity of the Y191 hydroxyl group (**Figure 1b**). Spectroscopic and reactivity data suggest that the conjugate base maintains a E/H232-H^+^-Y191^•^ triad to elevate the formal potential of the relay site so that it can effectively oxidize Fe(II) Cc. In both the Y191:E232 and Y191:H232 variants, pH dependencies indicate that proton loss from the E/H232-H^+^-Y191^•^, reduces the formal potential of the relay site, diminishing its ability to accept an electron from Fe(II) Cc [28].

**Figure 1.**
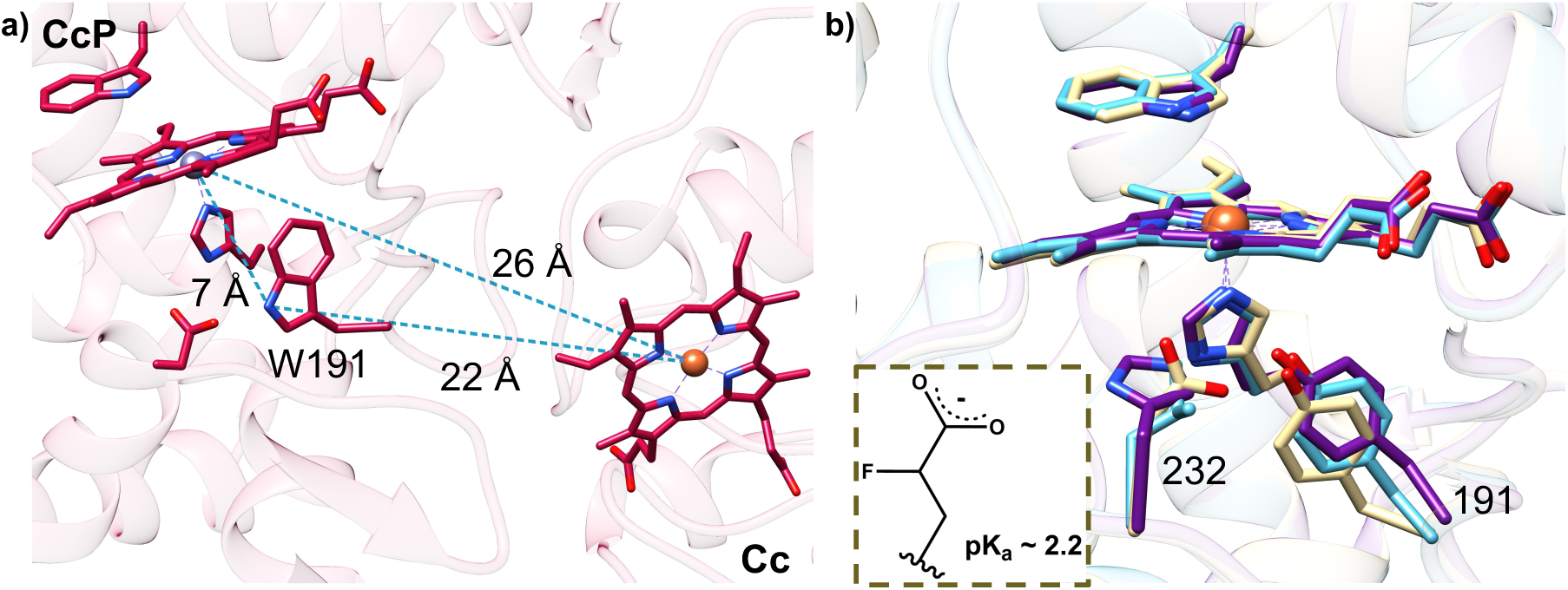
**a)** Distances between redox centers in the crystallographic CcP:Cc complex. **b)** Structural superpositions depict the Y191 environment in W191Y (or Y191, blue), W191Y:L232E (or Y191:E232, yellow), and W191Y:L232H (or Y191:H232, purple) CcP in complex with Cc. (PDB codes: 1u74, 5cih, 6p41, and 6p42). Inset: Molecular structure of 4-fluoro-glutamate.

Substitution of the CcP heme with zinc porphyrin (ZnCcP) allows light-activation of redox reactions with Cc [19,27,30–33] and thereby provides a useful system for further studying how the 191 relay site environment tunes ET (Scheme 2). For ZnCcP, 560 nm light excites zinc-porphyrin (ZnP) in CcP to the singlet state (^1^ZnP), which then rapidly decays to a long-lived triplet state (^3^ZnP), and in the absence of Fe(III) Cc relaxes back to the ground state with a rate constant, kD ∼100 s^-1^ [32–35]. In the presence of Fe(III) Cc, ^3^ZnP ejects an electron across the protein interface and reduces Cc, generating a charge separated species between ZnP^•+^ and Fe(II) Cc. Reduced Cc will then transfer an electron to CcP via hole-hopping through W191^•+^ with a similar ET mechanism as the native heme-peroxide system. The ability of a relay residue to accelerate net charge flux through a protein in part depends on the relay site formal potential relative to the donor and acceptor sites. In ZnCcP, a lower W191^•+^ potential relative to ZnP^•+^ localizes the hole closer to the donor, Cc (E° = 290 mV [36]), but also reduces the driving force to oxidize Cc. These factors play against each other in determining the net ET rate. The ZnCcP:Cc charge recombination reaction has been considered as a rapid hole exchange between ZnP^•+^ and W191^•+^ with the equilibrium lying toward ZnP^•+^ [37–39] i.e.

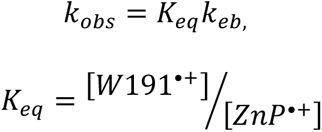

*K_eq_* ≪ 1, and *keb* being the back ET rate constant from Fe(II) Cc to CcP (Step 4 in Scheme 1 and Step 3 in Scheme 2 for FeCcP and ZnCcP respectively). However, there has not been a direct experimental measurement of *Keq*, a value that is challenging to assess given the high reactivity of the participating redox sites and low kinetic occupancy of the radical. In general, radical equilibration between multiple sites in proteins is difficult to characterize, but where these measurements have been achieved it is clear that the underlying factors determining reactivity are complex and difficult to predict [40–42].

**Scheme 2.**
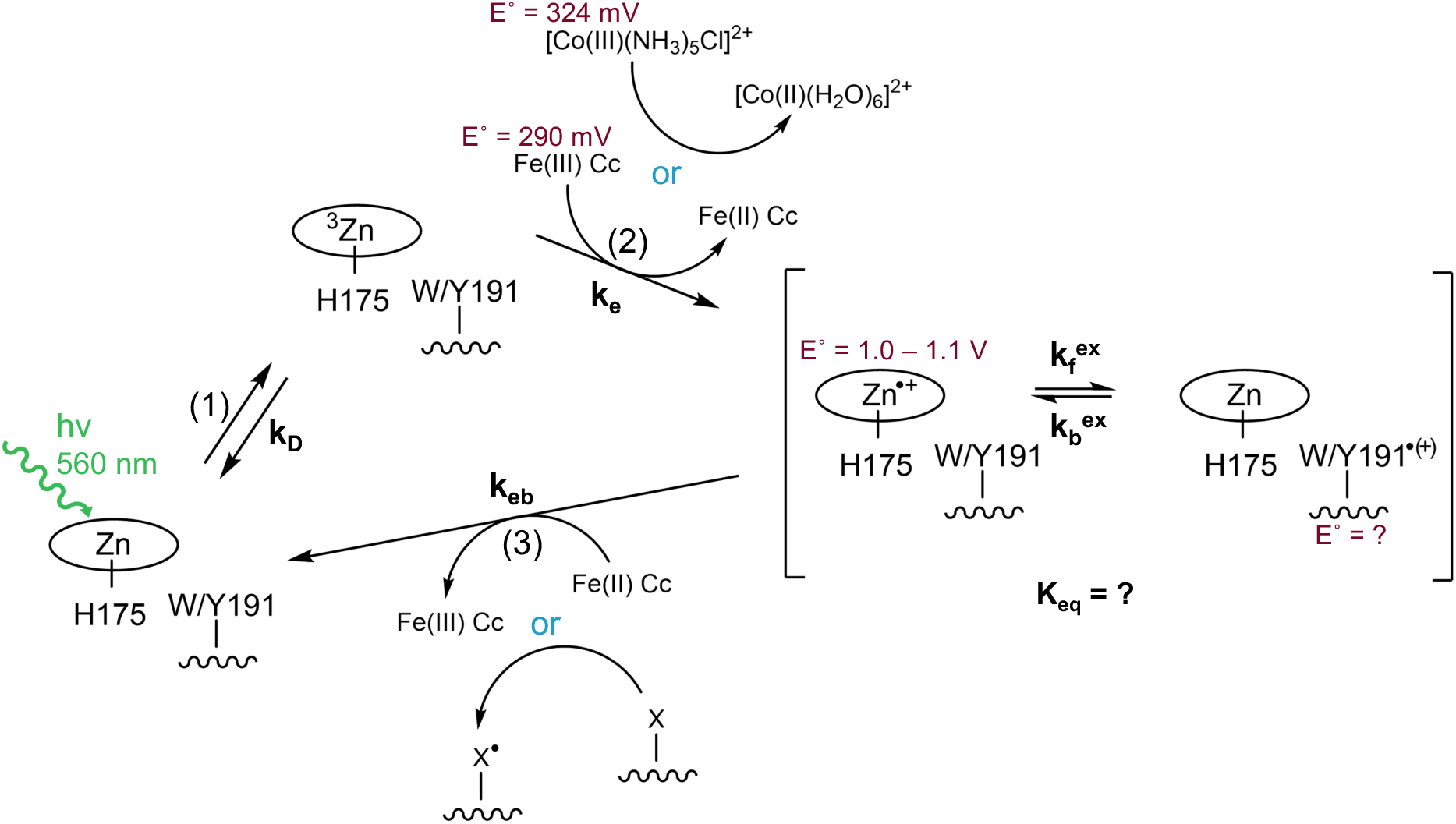
Photoinduced reaction scheme for ZnCcP

In efforts to leverage the well-studied CcP:Cc system to better understand the factors that allow relay sites to modulate protein ET, we have manipulated the p*K*a of the conjugate base by incorporating a fluorinated derivative of Glu, 4-fluoro-glutamate (F-Glu), and utilized a sacrificial electron acceptor ([Co(NH3)5Cl]^2+^) that allows stable generation of radical states after ZnCcP photoexcitation. Together the results underscore the importance of proton transfer at the relay site for both the peroxide-driven natural reaction of FeCcP and the photochemistry of ZnCcP. In the latter case, the nature of the base partner determines the ability of the radical to propagate elsewhere in the protein. These results provide insight into how minor perturbations to the protic environment of a redox active residue can substantially alter its reactivity. Turning on such reactivity can relocate high-potential sites to open up new channels for radical propagation. This information can potentially aid in the design of new protein-based catalysis and molecular systems designed to separate and mobilize charge.

## Results

### Cc oxidation assays of globally incorporated F-Glu FeCcP

Previous work suggested that a hydrogen bond provided by the neutral acid form of E232 to Y191 increased the reduction potential of the tyrosyl radical to a range where it could effectively oxidize Cc [28]. To test this mechanism, we sought to minimally perturb the E/H232-H^+^-Y191^•^ triad by lowering the p*K*a of the H-bonding residue such that it would become unprotonated at the pHs where ET activation was observed otherwise. The noncanonical amino acid 4-fluoro-glutamic acid is predicted to have p*K*a around 2 pH values lower than glutamic acid and will therefore be deprotonated at pH 5-8. F-Glu was globally incorporated into CcP using the endogenous translation machinery of an *E. coli* Glu auxotroph: PA-340 (*E. coli* Genetic Resource Center), in which CcP was expressed (**Table S1**). The endogenous translation system is unlikely to differentiate between Glu or F-Glu due to their high structural similarity. Hence, supplementation with F-Glu upon induction of CcP protein expression should result in at least partial F-Glu incorporation into CcP. To maximize F-Glu incorporation into the protein and limit toxicity associated with global incorporation, cells were initially grown to sufficient density in rich (Luria Broth) media, washed extensively with buffer and moved to a minimal media containing F-Glu, where expression was induced at 17 °C. Mass spectrometry (MS) data were collected of the purified F-Glu containing apo-CcP (**Figure S1**). Extracted ion chromatograms (XICs) indicated that ∼50 % of the ionized CcP tryptic peptide containing position 232 had F-Glu instead of Glu (**Table S2**). This is likely a lower bound on the incorporation level because F-Glu containing peptides are known to ionize less under ESI-MS positive mode than their Glu analogs [43].

In this approach, CcP can undergo global F-Glu substitution (F-EG) in any of its 20 endogenous Glu residues. Given the potentially complicating effects of global incorporation coupled with incomplete incorporation at the 232 site, we made reactivity comparisons among four variants: W191Y (or “Y191”), Y191 F-EG, Y191:L232E (or “Y191:E232”), and Y191:F-E232 F-EG. Rates of CcP Cpd I reduction by Fe(II) Cc were measured over multiple turnovers (Scheme 1) by monitoring Fe(II) Cc oxidation (550 – 540 nm) and progress of the ferryl Fe(IV)=O CcP (434 nm) intermediate in high ionic strength conditions (100 mM KP*i* buffer) over a range of pH 5 – 7 (**Figure 2**). Because these measurements were carried out under high ionic strength where the dissociation rates of the complex out compete the oxidation reactions, we expect the kinetics to reflect weighted averages of the reactivities provided by the heterogeneous F-Glu variants. Indeed, in all cases the initial oxidation rates are mono-phasic (**Figure 2 and Table S3)**. In multiple turnover conditions when Fe(II) Cc oxidation rates are slow relative to the WT CcP, Cpd I is regenerated from Cpd II (Fe(IV)=O, W/Y191^0^) by excess peroxide faster than Cpd II is oxidized by Cc [27,44]. Hence, the ferryl species (monitored at 434 nm, an isosbestic point for Fe(II)/Fe(III) Cc) increases as the concentration of Fe(II) Cc drops and peroxide is depleted. This gives rise to an initial fast phase where the majority of Fe(II) Cc is oxidized, followed by a slower phase where Cpd II is reduced to the ferric state by the remaining Fe(II) Cc [27,44]. The slow phase still involves participation of the radical at position 191 as the hole exchanges with Cpd II (Scheme 1). In the case of Y191:E232 and Y191:F-E232 F-EG CcP, the ferryl species grew in and decayed rapidly at low pH relative to the behavior of Y191 alone, commensurate with increased rates of Cc oxidation.

**Figure 2.**
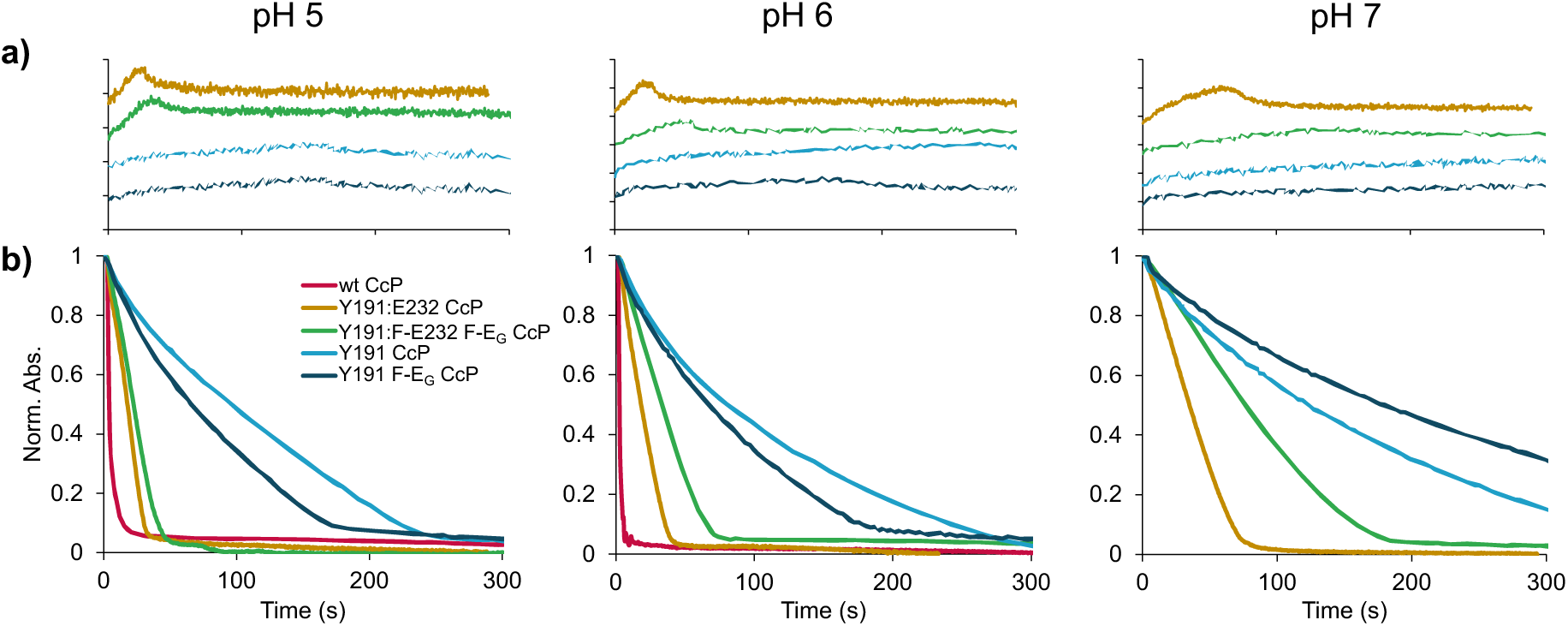
Cc oxidation rates of Y191 CcP variants under multiple turnover conditions (1 μM CcP, 30 μM Cc, 10 μM H2O2 in 100 mM KPi buffer). **a)** Time traces for Fe(IV)=O formation and decay monitored at 434 nm. Globally incorporated F-Glu CcP variants were monitored in comparison to unsubstituted counterparts in varying pH buffers. At higher pH, the ferryl CcP species in Y191:F-E232 F-EG is longer lived than in Y191:E232 CcP, similar to that of Y191 CcP. **b)** Time traces of Cc oxidation via CcP monitored at 550 – 540 nm. Both Y191:E232 and Y191:F-E232 F-EG exhibit pH dependencies in the rate of Cc oxidation.

In the early phase of this reaction Cpd I is replenished by peroxide faster than Fe(II) Cc oxidation [27,44] and thus to compare the activity of the different variants, the initial rates of Fe(II) Cc oxidation were obtained from fitting the kinetic traces to monoexponential decay functions (**Figure 3a**). Y191 and Y191 F-EG CcP showed virtually the same low activity and no pH dependence, indicating that F-E incorporation at non-232 sites had minimal effect on reactivity. At higher pH (pH 7), Y191:E232 and Y191:F-E232 F-EG exhibited slow Y191-like kinetics and Y191:E232 had an enhanced activity compared to Y191:F-E232 F-EG. However, at low pH (pH 5) the activities of Y191:E232 and Y191:F-E232 F-EG increased with Y191:E232 retaining greater activity than Y191:F-E232 F-EG (**Figure 3a**). Overall, the data indicates that E232 increases Fe(II) Cc oxidation rates, but in a pH dependent manner, such that rate enhancement diminishes as pH increased. The pH effect is most pronounced for Y191:F-E232 F-EG. A proton associated with E232 is then likely responsible for the rate enhancement.

**Figure 3.**
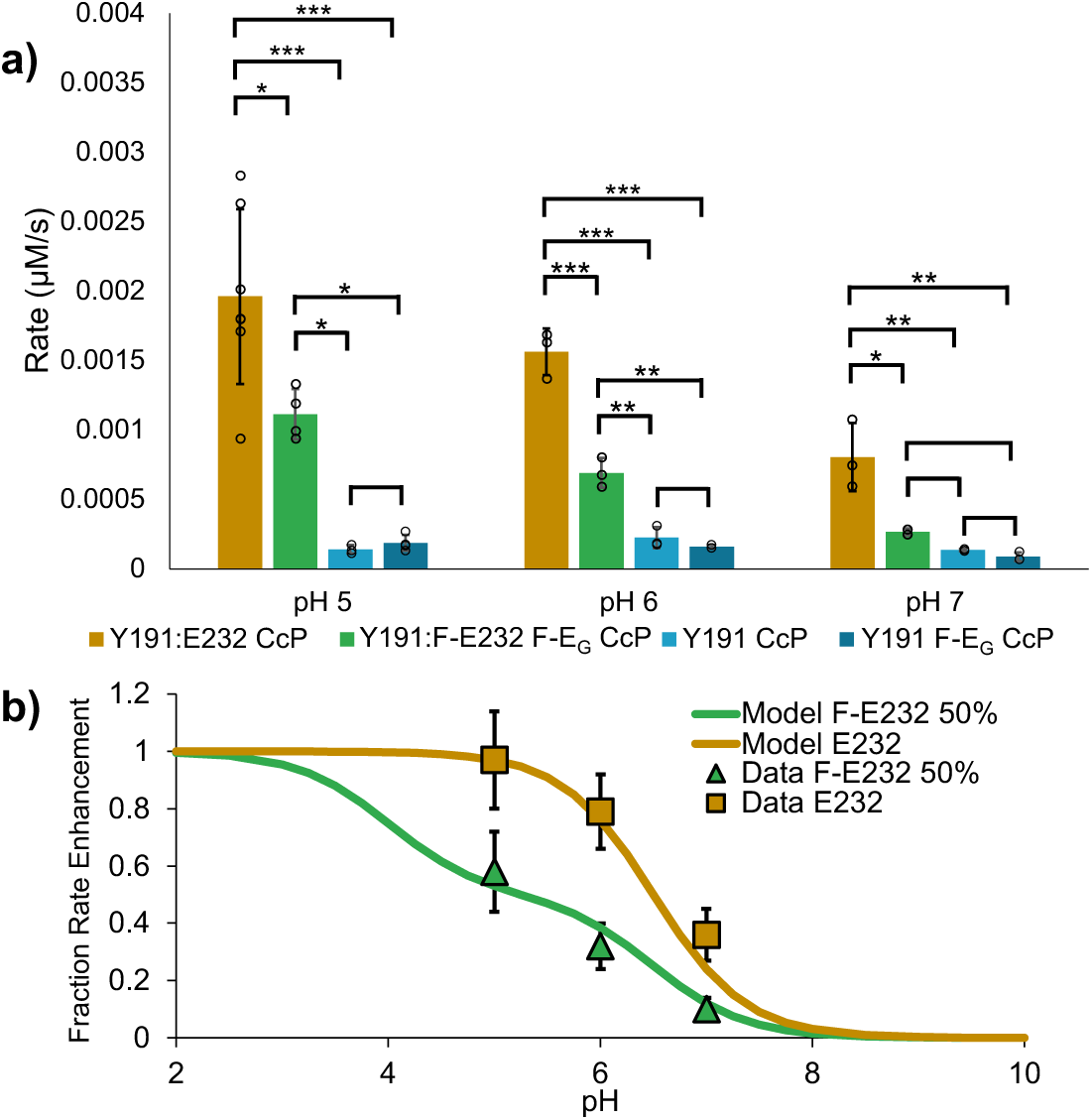
Initial rates of Cc oxidation by Y191 CcP variants. **a)** Y191 and Y191:F-E232 F-EG CcP lack any pH dependency and CcP activity is mostly lost. Y191:F-E232 F-EG was slower than Y191:E232 in all buffers regardless of pH and the Cc oxidation rate decreases to slower Y191-like kinetics at high pH for both variants. One-way ANOVA and post-hoc Tukey analysis was performed to evaluate statistical significance for each pH condition. Adjusted p-values are indicated where large statistical differences are observed (*** for p < 0.001, ** for 0.001 ≤ p < 0.01, and * for 0.01 ≤ p < 0.05). **b)** Comparison of a one and two proton ionization model to observed initial rates of Y191:E232 CcP and Y191:F-E232 F-EG CcP, respectively. Y191:F-E232 F-EG CcP rate assumes a 50:50 mixture of E232 and F-E232 in the sample.

Because global F-Glu incorporation does not greatly perturb CcP reactivity, we sought to better quantify the pH-dependent rate differences between Y191:E232 and Y191:F-E232 F-EG by considering the 50:50 mixture of Y191:F-E232 and Y191:E232. If the ability of E232 to accelerate ET from Cc depends on a hydrogen bond to Y191, the lower p*K*a of F-E232 may generate behavior similar to that of Y191:E232, but in a lower pH range. If we assumed a simple model wherein the Y191:F-E232 F-EG sample is a 50:50 mixture of CcP proteins that have fluorinated and unfluorinated E232 (as indicated by mass-spec), which differ in their p*K*a values (6.5 for E232 and 4.0 for F-E232), the equation below reasonably predicted the relative pH-dependency of each system. (**Figure 3b**).

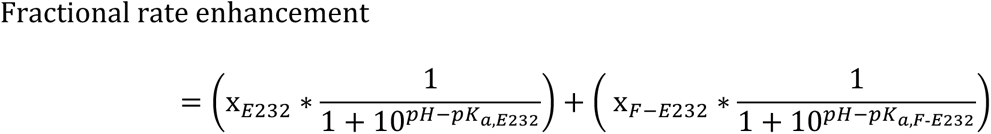

Where XE232 and XF-E232 correspond to the fraction of unfluorinated and fluorinated E232, respectively.

Difficulties in producing large amounts of heme-incorporated Y191:F-E232 F-EG and instability of the protein at low pH prevented a more extensive analysis; nonetheless, the rate behavior suggests that Y191:F-E232 F-EG CcP is unable to stabilize a proton in the F-E232-H^+^-Y191^•^ triad at elevated pHs (pH > 4) owing to its depressed p*K*a, and that the associated loss of a hydrogen bond to Y191 lowers its potential such that ET from Cc Fe(II) is reduced.

### PCET facilitates Y191 oxidation in the ZnCcP variants

The rate of charge recombination in the ZnCcP:Cc redox photocycles (Scheme 2) were measured with time-resolved spectroscopy across pH 6-8 for the Y191-substituted variants. In low ionic strength buffers, the CcP:Cc complex is stabilized against dissociation, but secondary site Cc binding can contribute to ET rates. Conversely, in high ionic strength buffers the active complex exchanges rapidly, but secondary site Cc binding is eliminated [34]. The smaller keb in Y191:E232 at high ionic strength indicates that complex dissociation does limit Cc oxidation on this time scale (**Figure 4**). Regardless of ionic strength, the Y191:E232 and Y191:H232 ZnCcP variants exhibited much greater pH dependencies compared to Y191 alone (**Figure 4, Table S4, S5**). For Y191:E232 at high ionic strength, keb increases 4-fold between pH 6 and 7, whereas keb for Y191:H232 increases as well at a higher pH of 7.3 (**Table S4**). This trend differs from that of the FeCcP peroxide reaction, where the Cc oxidation rates decrease at higher pH in both the E232 and H232 variants. For ZnCcP, the pH rate dependence also correlated with the relative p*K*a values of the adjacent Glu and His residues, but rates enhanced in the higher pH ranges. In this case, a proton acceptor may be needed to assist Tyr deprotonation during formation of Y191^•^ (**Figure 4**). Both high and low ionic strength conditions generate similar pH trends for Y191 and Y191:E232 (**Figure 4**). The pHs where rates increase for E232 and H232 (7 and 7.5-8, respectively) shifted higher than the p*K*as of the isolated residues (4.25 and 6.0 respectively), presumably owing to their protein environment, which includes proximity to both Tyr^•^ and ZnP^•+^. Notably, Tyr^•^ radicals are known to upshift p*K*a values of interacting bases in other systems [45].

**Figure 4.**
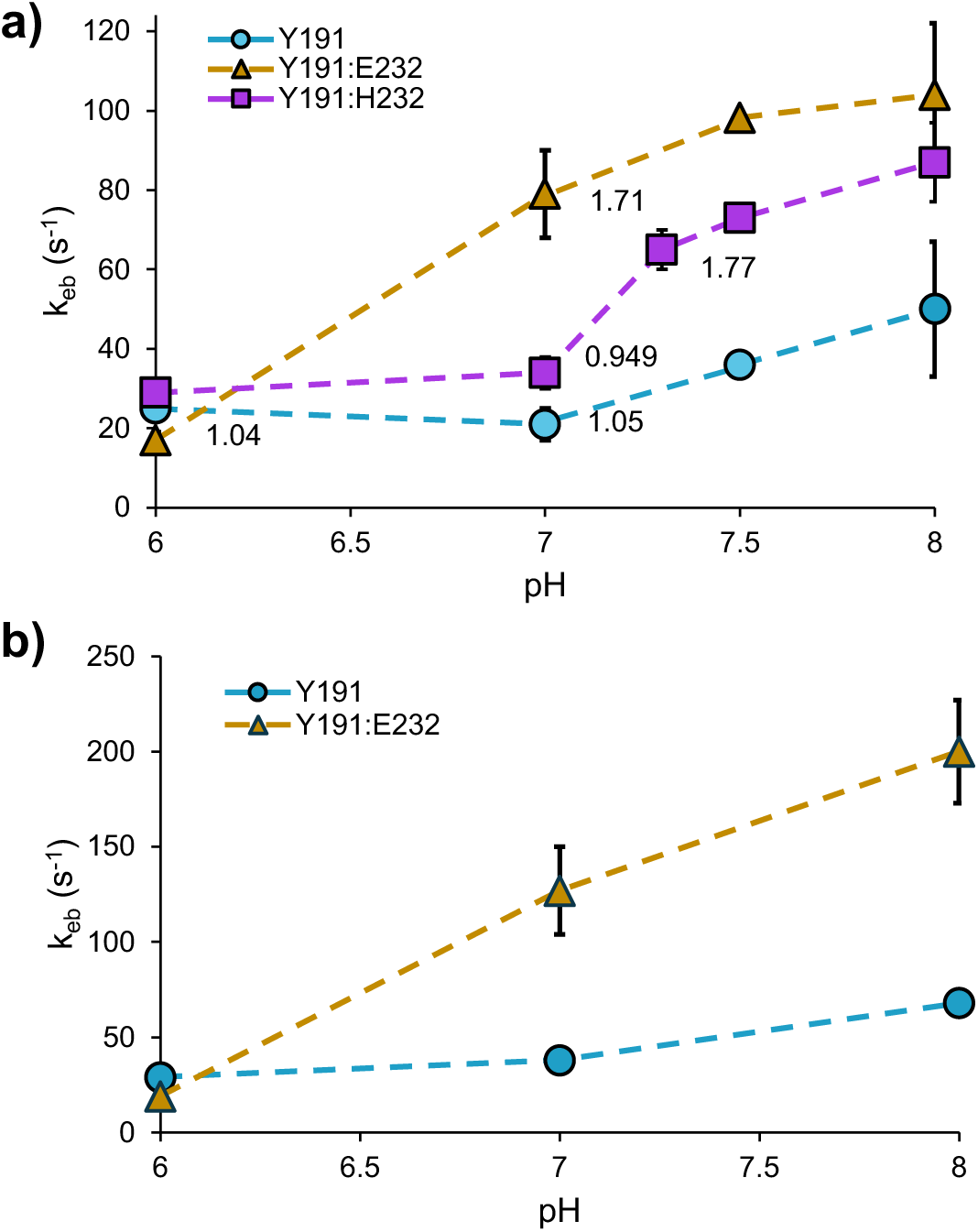
Back electron transfer rate constant (keb) as a function of pH for Y191 (blue), Y191:E232 (gold), and Y191:H232 (purple) CcP variants in complex with Cc in **a)** 100 mM KPi buffer and **b)** 10 mM KPi buffer. Kinetic solvent isotope effects (KSIE) are noted by corresponding data points when measured. keb has pH dependence for both the Y191:E232 and Y191:H232 variants and KSIEs track with increases in keb.

Consistent with a PCET mechanism, both the E232 and H232 variants showed a deuterium kinetic solvent isotope effect (KSIE), only at the pH values where rates increased. Y191:E232 gave a near 2-fold decrease in keb in the pD 7 deuterated buffer compared to the protonated pH 7 counterpart (**Table S6**). Similarly, Y191:H232 ZnCcP only exhibited a KSIE above pH 7.3, where keb also significantly increased. Notably, solvent deuterium substitution also modestly affects ionization properties (i.e. KW and p*K*a values) with the neutral point being ∼7.4 in D2O. This shift could affect the pH range over which the conjugate base ionizes and thereby potentially contribute to an apparent KSIE. However, we only observed a KSIE in the pH range where there was already rate acceleration, i.e. the conjugate base had ionized to act as a proton acceptor and hence the KSIE likely reflects the ability of the conjugate base to deprotonate Y191 in concert with oxidation by ZnP^•+^.

### Radical distributions in photo-oxidized ZnCcP

The difference in formal potential between ZnP^•+^ and W/Y191^•(+)^ is a critical parameter in the ability of ZnCcP to oxidize Cc [37], but because the recombination reaction of (ZnP/W191)^•+^ with Cc Fe(II) is faster than the reduction of Cc Fe(III) by the ^3^ZnP [38], (ZnP/W191)^•+^ is challenging to characterize. To address this issue, a sacrificial electron acceptor [Co(NH3)5Cl] Cl2 (CoN5) was used in place of Fe(III) Cc [46–49] (Scheme 2). When reduced, CoN5 decomposes into Co(H2O)62+, which diffuses and has a higher reduction potential than the precursor, and thus will not readily reduce ZnP^•+^ [49,50]. The transient absorption (TA) difference spectra of Y191 ZnCcP after photoexcitation with CoN5 revealed a ZnP^•+^ species at 690 nm that was stable for at least 10 minutes (**Figure 5a**). X-band continuous wave electron paramagnetic resonance (cwEPR) spectroscopy confirmed the presence of an organic radical in W191 ZnCcP and Y191 ZnCcP, whereas the G191 substitution produced a similar, but much lower amplitude cwEPR signal compared to W191 and Y191 (**Figure 5b**). The TA difference spectra indicates that this cwEPR signal arises in part from ZnP^•+^.

**Figure 5.**
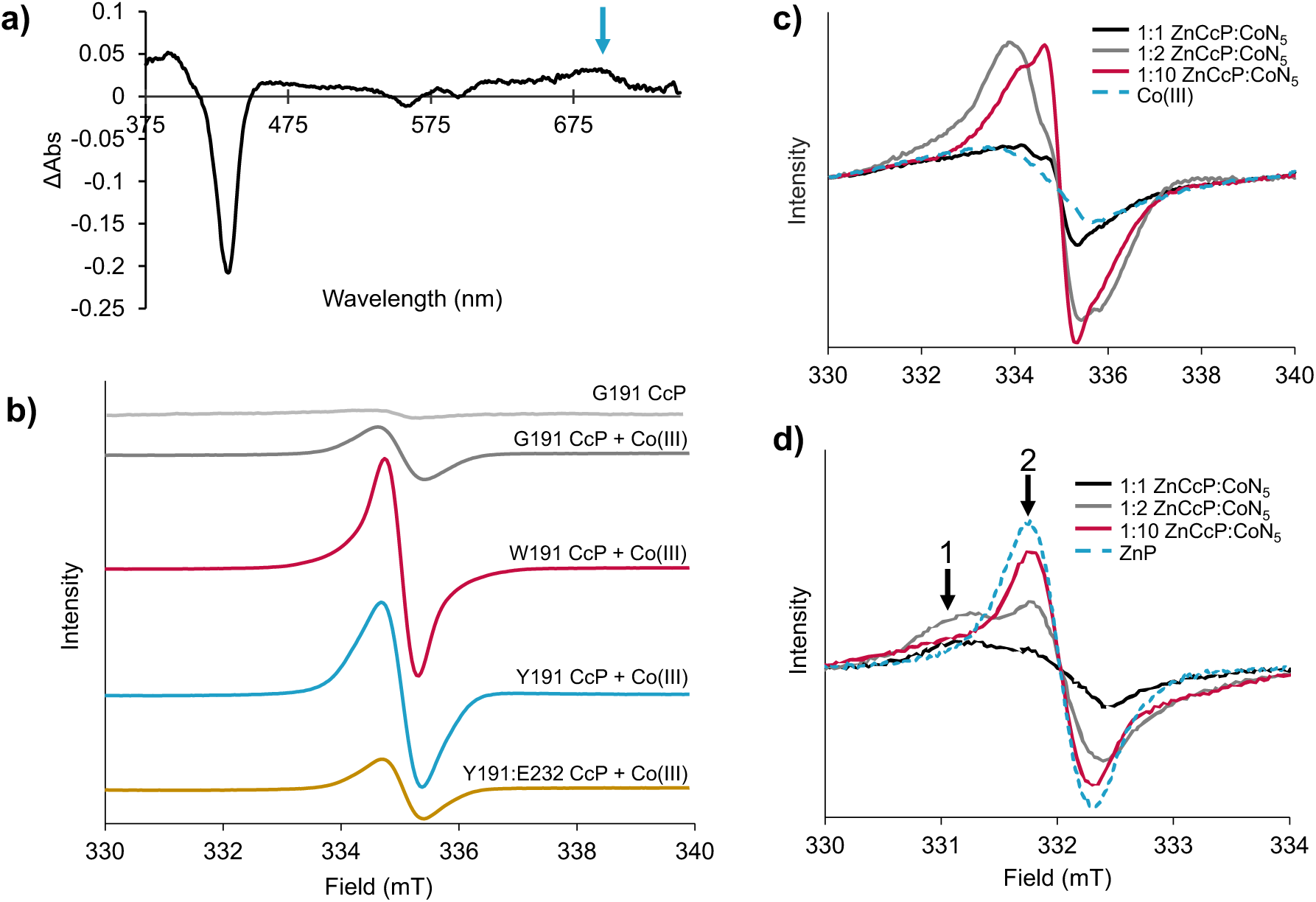
Characterization of of W191, G191, Y191, and Y191:E232 ZnCcP variants with CoN5. **a)** TA difference spectra of Y191 ZnCcP reveal ZnP^•+^ species (indicated by a blue arrow) at 690 nm. **b)** cwEPR at 10 K, pH 6 of ZnCcP mutants with 10-fold excess of CoN5 reveal organic radical formation present. In contrast, G191 CcP alone (light gray) has little to no signal in the spectra. The W191 containing wt CcP sample appears to have a slightly narrower signal than the other 191 variants indicative of a different radical species. cwEPR of wt ZnCcP at pH 6 with varying amounts of CoN5 at **c)** 10 K (0.4 mT modulation amplitude) and **d)** 110 K (0.5 mT modulation amplitude) reveal splitting of two radical species: the broad (1) and sharp (2) feature likely correlate with W191^•+^ and ZnP^•+^, respectively.

To better isolate the radical species contributing to the cwEPR spectra, the ratio between ZnCcP and CoN5 was varied. Initially, ZnCcP samples were measured at 10 K with 10-fold excess CoN5 to maximize yields of radical formation. Lowering the amount of CoN5 to stoichiometric ratios decreased the dominant resonance and revealed a previously obscured broad EPR signal (**Figure 5c**). Unlike the sharper fast-relaxing signal (T1 = 1.5 – 16 ms), the broad shoulder EPR transition relaxes slowly (T1 > 35 ms) and saturates with the microwave power used at 10 K. Power saturation studies carried out at 110 K revealed that the broad feature saturated at lower power than the sharper signal (**Table S7, Figure S2,3**). This difference, as well as the difference in relaxation times, indicates that the two features are unique radical species. With 40 dB attenuation (0.02 mW microwave power) at 110 K the signals became more separated and could be distinguished from each other (Labeled 1 and 2 in **Figure 5d**). Simulations of the spectra revealed that components 1 and 2 have unique apparent isotropic g-factors of 2.0033 ± 0.0001 and 2.0024 ± 0.0001, respectively (1:2 condition, **Table S8, Figures S4, S5**). The sharper, fast-relaxing feature matches well with the spectra of Zn protoporphyrin IX (g = 2.0023, **Table S8**), whereas component 1 is attributed to the oxidation of W191 by ZnP^•+^.

Stoichiometric amounts of CoN5 to ZnCcP gave rise to the highest relative signal of W191^•+^ compared to ZnP^•+^ (**Figure 5d**). Notably, no organic radicals directly form with light excitation of heme-containing CcP and CoN5 (**Figure S10**), thereby indicating that oxidation of ZnP precedes radical formation at W191. For the G191 variant at low CoN5 stoichiometry, cw-EPR, and measurements of Tm and T1 at 60 K and 110 K indicated a high proportion of the fast-relaxing ZnP^•+^ (**Table S9**). The lack of substantial radical propagation in G191 ZnCcP, differs from the heme-peroxide reaction, wherein the G191 FeCcP:Cc primarily produces a peripheral Tyr^•^ radical upon reaction with peroxide [24]. This difference in reactivity between FeCcP and ZnCcP G191 suggests that the peroxide-derived Cpd I more readily oxidizes peripheral sites because it has a higher potential than ZnP^•+^. In all cases, there was no dipolar line broadening of the ZnP^•+^ signal, which appeared the same among the variants, including G191. Interestingly, ZnP^•+^ was less stable compared to W191^•+^, and overnight at 293 K, ZnP^•+^ decayed, leaving only the W191^•+^ signal, whose linewidth decreased slightly (**Figure S4, Table S10**). The W191^•+^ signal also broadens slightly when CoN5 to CcP ratios increase and more ZnP^•+^ is present at 110 K (**Figure S5a-c**, **Table S8**). ZnP and W191 are separated by 7-8 Å (**Figure 1a**). Simulations of dipolar line-broadening of the 110 K data predict that bi-radicals at this distance would induce a peak-to-peak linewidth in the range of 4-5 mT, much larger than what is observed (**Figure S5d,e**). Therefore, bi-radicals at this separation do not contribute appreciably to these spectra.

Photoreduction of the Y191-substituted CcP variants with CoN5 at stoichiometric ratios were studied by cwEPR by in situ steady-state irradiation at room temperature. This approach produced cwEPR signals characteristic of ZnP^•+^ with a smaller contribution from a broad shoulder near the ZnP^•+^ peak, which may be from tyrosyl or another organic radical in the sample (X^•^, **Figure 6b-d**). Furthermore, the spectra were pH dependent, with the ZnP^•+^ signal dissipating at higher pH, but the broader, low-field signal persisting. The pH dependence also varied among the Y191 variants, with loss of the ZnP^•+^ signal at pH 8 for the Y191 variant, pH 7 for Y191:E232 and pH 6 for Y191:H232 (**Figure 6, Figures S6-S9, Table S11**). These results relate to the ET kinetics of the ZnCcP:Cc complex, where higher pH values (7-8), increase the ET rate from Cc owing to greater yield of radical character at the relay site. For Y191:E232, higher pH deprotonates E232, which then accepts a proton from Y191 to form the tyrosyl radical; for the Y191 without a conjugate base, yet higher pH is required for solvent to likely assist in Y191 deprotonation; for Y191:H232 the EPR spectra surprisingly showed no pH dependence with the broad, low-field peak present at pHs 6-8 and little indication of ZnP^•+^. One might expect H232 to deprotonate at even higher pH than E232, but the close proximity of ZnP^•+^ and structure of the site may favor the interaction of the neutral imidazole with the Y191 hydroxyl and thereby poise the Y191 proton for transfer across the pH range.

**Figure 6.**
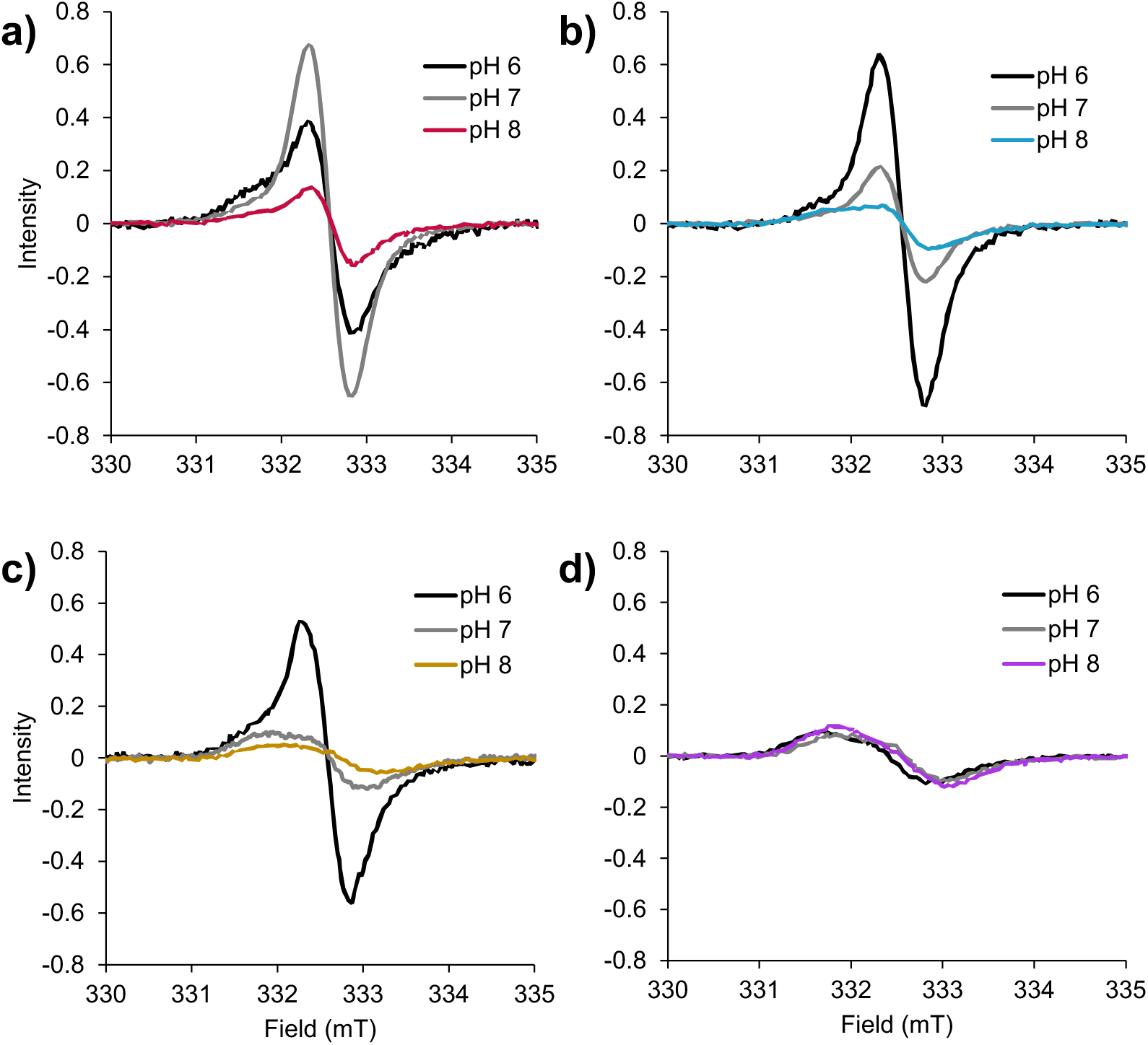
cwEPR spectra recorded at 293 K (0.4 mT modulation amplitude) of **a)** W191, **b)** Y191, **c)** Y191:E232, and **d)** Y191:H232 variants with one-molar equivalent of CoN5. Y191 and Y191:E232 ZnCcP variants exhibit pH-dependent behaviors of their cw spectra that track similarly as their ET rates in Figure 4 where X^•^ is more prevalent in high pH conditions. Y191:H232 ZnCcP differs from the others in that the low-field radical is the only species present at all pH conditions.

Pulsed EPR techniques were carried out to further characterize the radical species generated in each ZnCcP variant by CoN5 quenching. ^1^H Davies electron nuclear double resonance (ENDOR) spectra of W191 ZnCcP at high CoN5 stoichiometry (10:1) and low temperature 10 K, revealed hyperfine coupling interactions characteristic of free ZnP^•+^ in solution (**Figure S10**). There are also broad hyperfine interactions at ∼15 MHz that resemble those of W191^•+^ from CcP Cpd I [22]. To better isolate the hyperfine interactions of the non-ZnP^+•^ radical, X^•^, ^1^H Davies ENDOR was carried out at low CoN5, 110 K, pH 6, with the ENDOR resonance centered on the low field peak (**Figure 7a, S11**). The ZnP^•+^ control showed few hyperfine interactions at this field, but the W191 ZnCcP gave broad, considerably shifted peaks, characteristic of W191^•+^, which are consistent with simulated data from published parameters for the W191^•+^ species of CcP Cpd I [51] (**Figure S12**). Note that in Cpd I, W191^•+^ is weakly coupled to the S=1 ferryl heme, which is not the case here [22,25,26]. Y191 and Y191:E232 produced similar ENDOR spectra to each other under these conditions, but that of Y191:H232 contains additional shoulder peaks indicative of X^•^ (**Figure 7a**). At pH 8, the Y191:E232 spectrum resembles more closely the Y191:H232 spectrum, as it also contains shoulder peaks. The G191 ENDOR spectrum also contains some peaks similar to ZnP^•+^, but no broad shoulders (**Figure 7a**).

**Figure 7.**
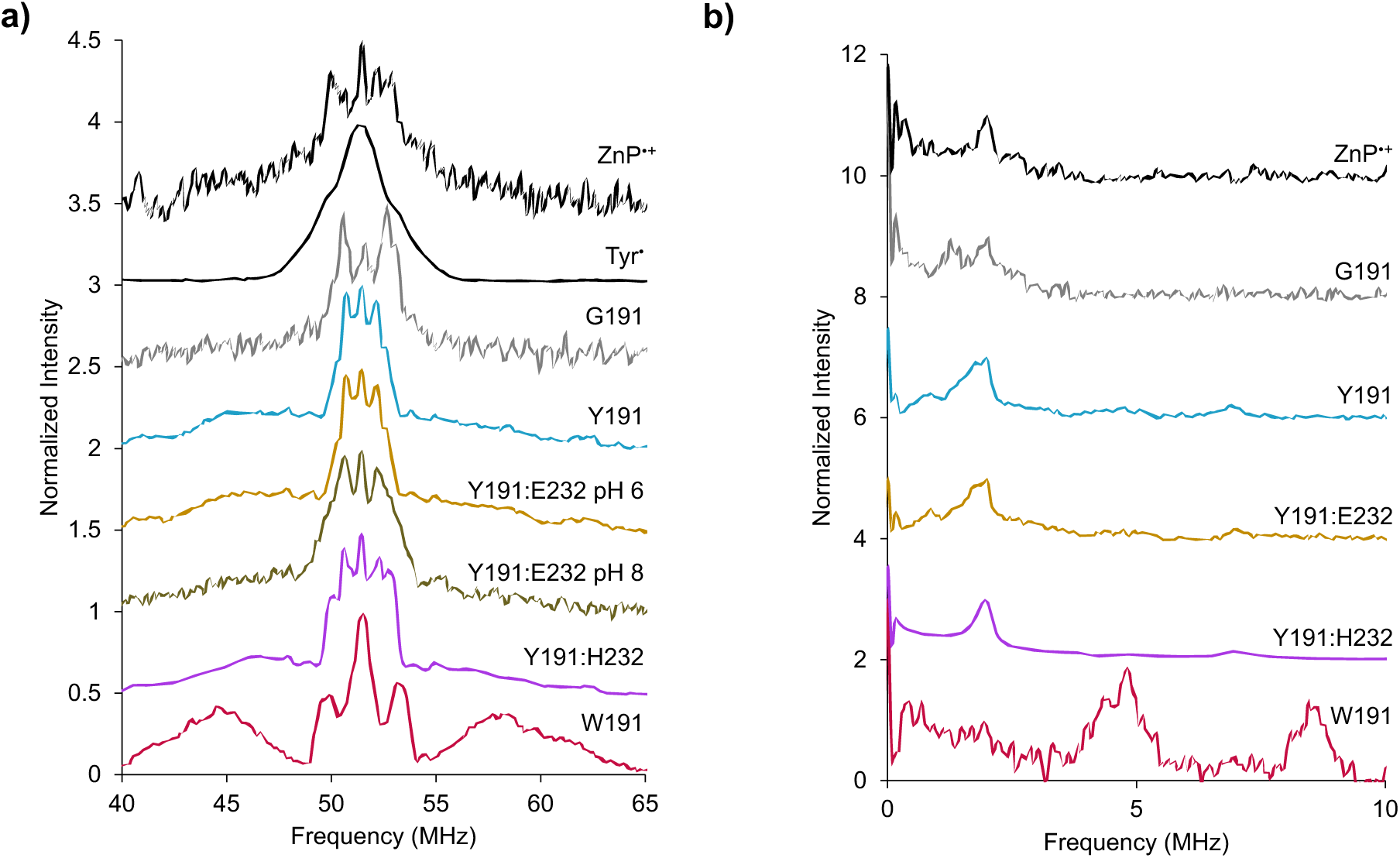
Hyperfine spectroscopy of ZnCcP radical species. **a)** ^1^H ENDOR spectra of free ZnP^•+^and Tyr^•^ and of ZnCcP variants at 110 K, pH 6 (and pH 8 for Y191:E232) with equimolar CoN5 normalized to the largest peak. The EPR observation frequency at 34 GHz was chosen to emphasize proton coupling to the low-field organic radical (**Figure S11**). Central peaks representing the proton Larmor frequency are present in spectra of all ZnCcP species. Signals from hyperfine-shifted Trp^•+^ protons are also present in W191 ZnCcP at 15 MHz whereas Y191 variants contain a broad peak at 8 MHz. G191 also exhibits some shoulder, but signal-to-noise is low. Tyr^•^ signal is shown for comparison. **b)** ^14^N ESEEM spectra at 60 K, pH 6 also centered on the low-field radical transition of free ZnP^•+^ and ZnCcP variants with equimolar CoN5 normalized to the 1-2 MHz peak. A sharp peak at 2 MHz is attributed to ^14^N-coupling in the porphyrin. Spectra of the W191 and Y191 variants both include additional peaks at 5 and 7-8 MHz, likely due in part to hyperfine interactions with indole nitrogen in Trp residues.

14N electron spin echo envelope modulation (ESEEM) spectroscopy was utilized to further characterize X^•^ (**Figure 7b**). Resonances in free ZnP^•+^ and G191 ZnCcP at 2 MHz arise from the coupling of the ZnP radical to nitrogen in the porphyrin pyrrole rings. This feature is also observed in the Y191 ZnCcP variants where ZnP^•+^ is still present in appreciable amounts. In addition to this interaction, both the W191 and Y191 variants also exhibit a pair of peaks with resonances located around 5 MHz and 7-8 MHz. With these coupling intensities and frequencies, it seems unlikely that even in the 191 variants X^•^ derives from only a tyrosyl radical, which usually has no nitrogen hyperfine coupling [52–54] or very weak (<1 MHz) nitrogen couplings from residues in the vicinity of the tyrosyl radical [55]. This nitrogen coupling behavior indicates that a migrated tryptophanyl radical may be at least partially responsible for these signals (**Figure 7b**). The positions of ESEEM signals were generally insensitive to temperature but accentuated at 60 K compared to 110 K (**Figures S10**). Simulation of the W191 ESEEM-derived hyperfine parameters (**Figure S13**) matched reasonably well with the ENDOR results and those calculated previously [51]. For the Y191 variant ESEEM spectra, only the ∼2 MHz peak could be accurately simulated, and this feature matched that found in the spectrum of ZnP^•+^. However, the ∼5 and ∼7 MHz features are unique to the Y191 variants and argue in favor of some other nitrogen-containing radical, possibly a peripheral Trp^•^ (**Figure 7b, S14**). Thus, with CoN5 replacing Cc and preventing recombination, the primary ZnP^•+^ radical partially migrated through Y191^•^ to peripheral sites, potentially Trp residues. Radical migration depended on 1) first forming Y191^•^, which is facilitated by Y191 deprotonation and 2) maintaining a suitably high Y191^•^ potential to oxidize peripheral sites, which is facilitated by hydrogen bonding from the deprotonating residue. Thus, proton environment modulates Y191 reactivity in several ways, producing different pH dependencies for different variants: the Y191 variant activates the relay site to migrate the radical at pH 8, Y191:E232 does so at pH 7 or higher, and Y191:H232 does so at pH 6 or higher.

### QM/MM -MD calculations

QM/MM structure optimizations followed by spin density analysis along QM/MM MD trajectories revealed distinct radical localization patterns in the W191 and Y191 ZnCcP variants (**Figure 8, S15-S16**). In (ZnP/W191)^•+^, alternating radical character between ZnP^•+^ and Trp^•+^ was observed over the MD trajectory, consistent with EPR features for both radicals when the CoN5 stoichiometry was low. The calculations suggest that Keq lies toward Trp^•+^, with the Trp^•+^/ZnP^+^ratio ∼ 1.8, which agrees with the distribution found by EPR spectroscopy, averaged over the pH range of measurements (**Table S11**, **Figure S17**). It is interesting that the QM/MM simulations here yield different results from previous work [38,51], which found greater delocalization of the hole over (ZnP/Trp)^•+^ and hole occupancy favoring ZnP^•+^. We consider two possible reasons for the discrepancy. First, whereas both studies use DFT to calculate spin density in the QM region, the wB97x functional employed here is a range-separated hybrid that proves more accurate for spin densities than B3LYP (additional discussion in the SI). Second, because the prior work focused on the larger question of identifying the residues that participate in the long-range ET process, longer less accurate MD trajectories were employed [38], whereas the present study is focused on the question of hole localization and the effect of proton transfer on the relay residue and employs shorter but more accurate QM/MM MD trajectories.

**Figure 8.**
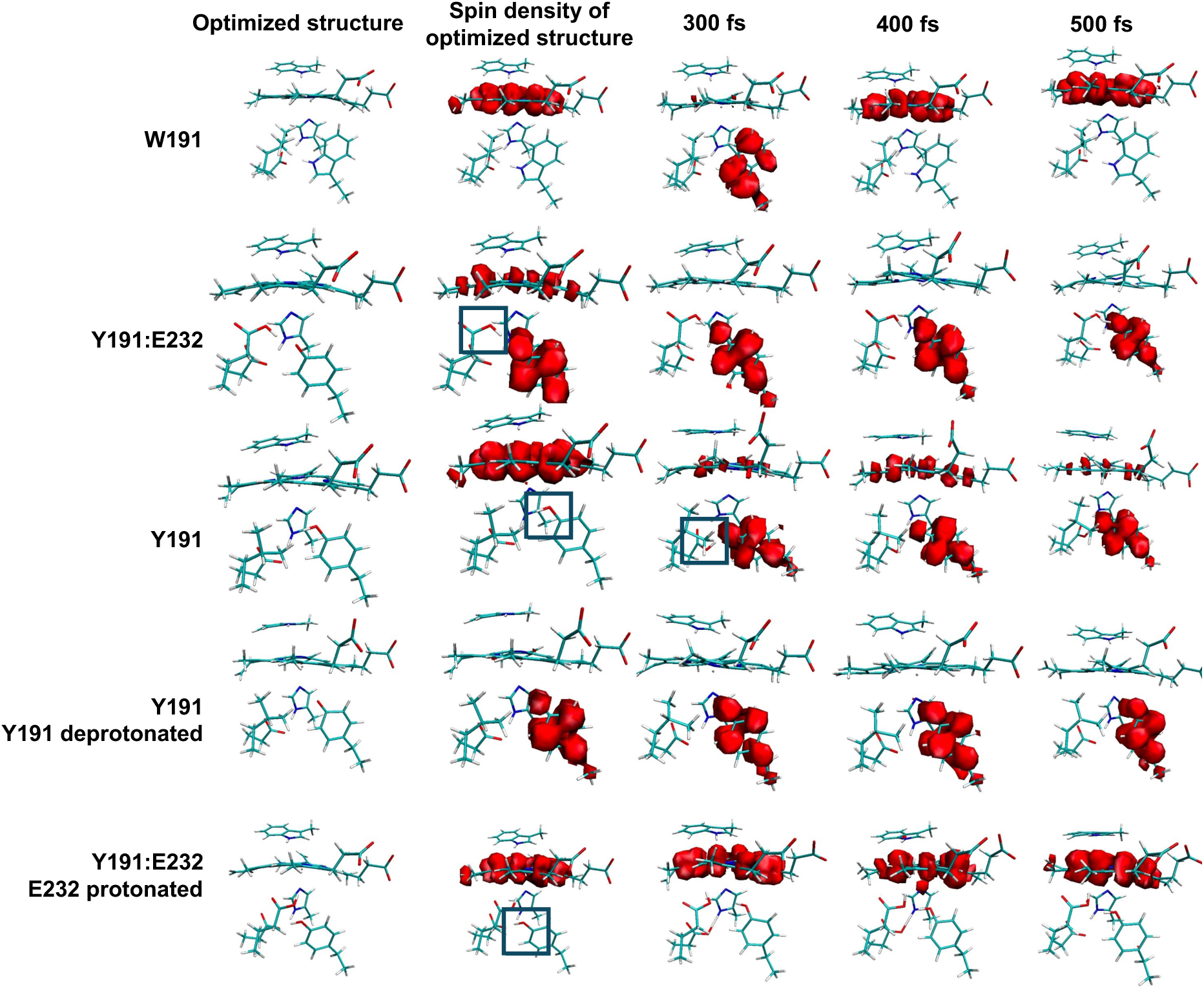
Spin density obtained from DFT with the wB97x functional as a function of time along the QM/MM MD trajectory for the different variants characterized experimentally. Three snapshots drawn from a 1ps long trajectory (additional structures in SI **Figures S17-S21**) highlight the key differences in behavior of the three variants and the importance of PCET in enabling hole transfer between ZnP and W191/Y191. E232 deprotonation facilitates proton transfer and radical formation at Y191. Boxes indicate location of proton in the optimized structure.

For Y191:E232 with E232 deprotonated, the simulations indicated proton transfer from Y191 to E232 during structural optimization, accompanied by radical localization on Y191^•^ (**Figure 8**, Row 2), thereby supporting a PCET mechanism that is consistent with the experimentally observed solvent isotope effects. When Y191:E232 was modeled with protonated (neutral) E232, spin density localized exclusively on ZnP^•+^, (**Figure 8**, Row 5) underscoring the role of proton transfer in facilitating Tyr^•^ formation and corroborating the pH-dependent rate studies. If both Y191 and E232 were deprotonated (**Figure 8**, Row 4), the radical localized to Y191^•^ exclusively, whereas when proton transfer is possible (**Figure 8**, Row 2), we find some spin density on ZnP^•+^. For the Y191 variant, (ZnP/Y191)^•(+)^, when the Y191 proton transfers to D235, the spin density again predominantly localizes on Tyr but partial ZnP^•+^ radical character again appears when a hydrogen bond is formed between D235 and Y191^•^ during the QM/MM trajectory (**Figure 8**, Row 3). Thus, like in the case of E232, a hydrogen bond back to Y191^•^ drives radical character partially back to ZnP.

## Discussion

The well-studied CcP:Cc system provides a sound template to explore radical propagation reactions in proteins, which are notoriously difficult to study given the challenges in synchronizing these reactions and the high reactivity of the species involved. The W191^•+^ hole-hopping site is necessary for effective long-range ET in the CcP:Cc system, but it is also sensitive to immediate environment. For FeCcP Y191, an ionizable E/H232 residue activates the Y191 relay site, but somewhat surprisingly, only at low pH, where these residues are expected to be protonated [28]. This result could be explained by the ability of E/H232 to provide a hydrogen bond to the Y191^•^ radical and thereby facilitate its ability to oxidize Cc. With increasing pH, the E/H232-H^+^-Y191^•^ triad deprotonates and the subsequent lowering of the Y191^•^ reduction potential reduces Cc oxidation rates.

Our goal was to test this hypothesis by further lowering the p*K*a of the conjugate base in the absence of any other structural effects and examine the effect of pH on rate enhancement. Incorporation of F-E232 into CcP was challenging because F-Glu is isosteric with Glu and hence substitution is difficult to achieve by genetic-code expansion methods. Semi-synthesis approaches [56] are also complicated because of the need to associate a cofactor with CcP. We thus applied an auxotrophic supplementation strategy that was reasonably effective at incorporating the modified residue (∼50 %), despite the accompanying complication of global F-Glu substitution throughout the proteome. Nonetheless, the loss of rate enhancement at lower pHs by F-Glu substitution indeed supports the assumed mechanism (**Figure 3**). Based on the predicted p*K*a of F-Glu, we would expect the Y191:F-E232 F-EG CcP, Fe(II) Cc oxidation rates to increase even further at lower pHs (3-4), and because F-E232 is a better proton donor than E232, the hydrogen bond donated to Y191^•^ may be more effective at increasing the Y191^•^ redox potential than with E232. Unfortunately, it was challenging to measure ET at low pHs due to protein stability. Nonetheless, the F-Glu behavior strongly implicates the importance of single proton for enhancing Y191^•^ reactivity, because structurally, F-Glu is nearly identical to Glu.

Remarkably, rate enhancement in the peroxide-activated FeCcP:Cc system shows an opposite pH trend compared to the photo-activated ZnCcP:Cc system, wherein higher pH increases, not decreases Cc oxidation rates (**Figure 4, Table S4**). This finding reflects how changes in reduction potentials can perturb mechanisms of radical transfer. For FeCcP, a hydrogen bond to Y191^•^ from E/H232 provides a high-enough reduction potential to oxidize Fe(II) Cc. However, for ZnCcP, the preceding oxidation of Y191 by ZnP^•+^ is likely the pH-sensitive reaction (**Figure 9a**). PCET from Y191 to deprotonated E/H232 (or solvent) facilitates Y191 oxidation by ZnP^•+^. A similar effect is observed in PSII where the oxidations of both TyrD and TyrZ are coupled to proton transfers to hydrogen-bonded His residues; when the His residues are protonated at low pH, the Tyr residues are nonfunctional as relay sites [54,57]. Moreover, function of TyrZ could be restored by solvent or high pH when the hydrogen-bonded base was removed, indicative of TyrZ deprotonation to solvent upon oxidation [12–14]. ZnCcP reactivity implies that Cpd I in the peroxide CcP reaction has a higher reduction potential relative to Y191 than ZnP^•+^ in the photochemical system because with FeCcP, the pH sensitive reaction involves the latter step, i.e. oxidation of Fe(II) Cc by the already formed Y191^•^ (**Figure 9b**). Notably, even if the lower reduction potential of ZnP^•+^ compared to Cpd I requires PCET to activate Y191^•^ in the former case, a subsequent hydrogen bond back from the conjugate base to Y191^•^ may still facilitate the oxidation of Cc Fe(II). For ZnCcP, the pH range where rate enhancement manifests is somewhat higher than that predicted by the p*K*a of free Glu. However, it is not uncommon for the p*K*a of residues within the protein interior to be perturbed 5-6 pH values higher than their solution p*K*as with Glu residues having p*K*a increases from 4 to around 7 [58,59].

**Figure 9.**
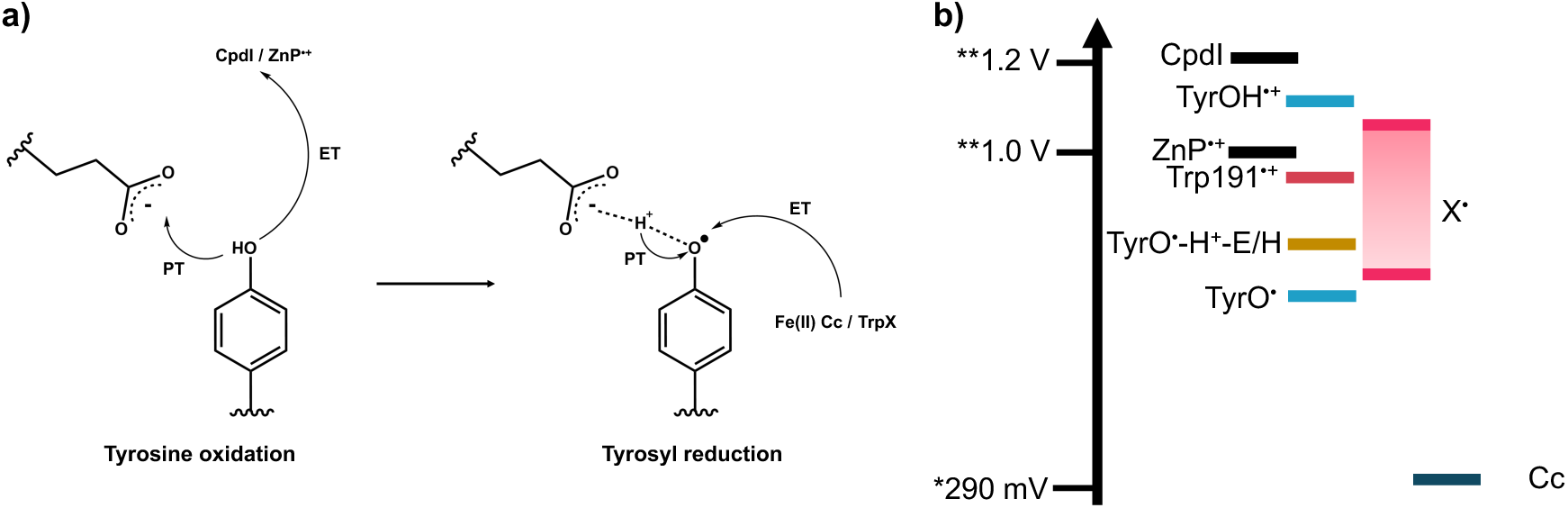
**a)** PCET steps in Cc oxidation by CcP. In ZnCcP, the oxidation of Y191 becomes rate limiting because the TyrOH^•+^ potential is too high relative to that of ZnP^•+^ for the Tyr to be oxidized in the absence of PCET. **b)** Proposed potential energy diagram for oxidizable species in CcP:Cc. *Taken from [36], ** Taken from [38].

The difference in reduction potentials between reactive sites (in this case ZnP^•+^ and W/Y191^•(+)^) is a critical parameter in determining the efficacy of ET relay. Past computational studies have estimated Keq < 1 [37–39], which effectively downweighs the otherwise large rate constant for ET between Fe(II) Cc and Trp^•+^ (> 10^8^ sec^-1^) [39]. Use of CoN5 as an oxidative quencher to photoexcited ZnP stabilized the protein radical against recombination with the external reductant allowing for direct assessment of the relative potentials between the two sites. No radical species were observed when FeCcP underwent the same irradiation protocol (**Figure S10a**), thereby indicating that W191^•+^ formation depends on ZnP^•+^ and the cation radical distributes between the two sites, favoring Trp^•+^. Both the experimental fits to the EPR data (**Table S11**) and the QM/MM trajectories give 1:1 – 2:1 ratios of Trp^•+^/ZnP^•+^ after ZnCcP oxidation, hence placing the Trp^•+^ formal potential between -1.4 and 27 mV apart from that of ZnP^•+^ (1.0 - 1.1 V) [38], which is not a very significant difference (**Figure 9b**). With kobs ∼ 10^6^ s^-1^ [60] and Keq > 1, the ET rate between ZnP^•+^ and W191^•+^ must be considerably slower than expected. Indeed, the ZnP^•+^ and W191^•+^ resonances differ by ∼0.1-0.5 mT (Δν ∼ 3-14 MHz) and kfex the ET rate corresponding to:

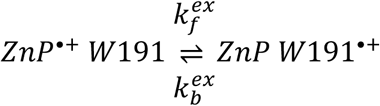

must be less than this value, i.e. < 1×10^7^ s^-1^. Despite their relatively close proximity, ZnP and W191 are not effectively coupled, presumably owing to the nature of the participating and intervening molecular orbitals. Thus, the protein scaffold may modulate donor-acceptor coupling, perhaps through orientation effects.

For oxidative quenching of Y191 ZnCcP with CoN5, a low-field broadened peak also appears in the EPR spectrum along with the characteristic ZnP^•+^ signal; and its population depends differently on pH among the 232 variants. ESEEM suggests ^14^N hyperfine coupling to this transition, and thus some contribution from a tryptophanyl radical, X^•^, is most likely (**Figure 7b**). For Y191:H232, X^•^ is dominant at pH 6-8, with little contribution from ZnP^•+^, for Y191:E232, this is the case for pH 7 and above, and for Y191 alone, X^•^ begins to dominate at pH 8. In all three cases, the spectral transition is associated with loss of ZnP^•+^ because of its ability to oxidize the 191 position. Consistent with the ZnCcP:Cc oxidation experiments, a hydrogen bond acceptor (E/H232) close to Y191, facilitates PCET from Y191 to ZnP^•+^, but the pH range where Y191 activates, depends on the 232 conjugate base. Y191:H232 propagates the radical most effectively at low pH, where the H232 should remain protonated (p*K*a ∼ 7.3 as indicated by the ZnCcP:Cc kinetics) and unable to assist in Y191 deprotonation. However, the apparent H232 p*K*a may be different in the absence of Cc and presence of CoN5 and under these equilibrium conditions. H232 is 3.4 Å away from H175 which is coordinated to the Zn atom in ZnP, whereas E232 is more than 4 Å away from H175. The close proximity of ZnP^•+^ may also disfavor the positively charged imidazole cation. Thus, it is possible that under the CoN5 conditions, the local environment maintains a more standard p*K*a for H232 (i.e. p*K*a = 6). When Y191 oxidizes, and the hydroxyl proton transfers to the base, the radical at least partially migrates from Y191^•^ to X^•^. The ENDOR spectra of Y191:E232 resembles that of Y191 at pH 6, but at pH 8, additional hyperfine shoulder peaks arise in Y191:E232, much like those seen for Y191:H232 (**Figure 7a**). Thus, a hydrogen bond from the conjugate base back to Y191^•^ may increase the Y191^•^ potential sufficiently to at least partially oxidize these peripheral sites, whose identities remain undetermined.

The QM/MM calculations further support this interpretation. In the optimized structure, deprotonated E232 accepted a proton from Y191, which coincided with Y191^•^ formation. Some radical character localized to ZnP^•+^ at the beginning of the simulation. No radicals localized on Y191 if E232 began the simulation protonated, unable to accept a proton from Y191, and no radical character localizes to ZnP, if E232 and Y191^•^ do not share a hydrogen bond. The Y191 alone simulation also emphasized the importance of proton transfer in stabilizing Y191^•^ radical character. With no E232 present, the Y191 proton transfers instead to D235 and radical character localizes to Y191^•^, with some radical character again on ZnP^•+^ when protonated D235 hydrogen bonds back to Y191^•^. In the actual protein it is unlikely that D235 hydrogen bonds to Y191^•^, because without E232, the ET reaction is slow and pH dependencies are different, but the behavior of D235 in the simulation demonstrates that a general base is necessary to deprotonate Y191, and that the resulting hydrogen bond to Y191^•^ facilitates some radical distribution back to ZnP^•+^, presumably by raising the formal potential of Y191^•^.

In the experimental setting, the radical on Y191^•^ migrates away from ZnP to other sites in CcP. Regardless of the identity of the migrated radical in the Y191 variants, ZnP^•+^ does not oxidize Y191 as easily as W191 unless oxidation couples to Y191 deprotonation; a PCET effect could then arise from the first ET step of Y191 to ZnP^•+^, (**Figure 9a**). Both the peroxide dependent reaction [28] and the ZnCcP photochemical oxidation of Cc show a modest KSIE under conditions where the relay site is activated and rates are enhanced. These KSIEs are similar to those for the PT-coupled reduction of the TyrD^•^ in PSII, but unlike those of model β-hairpin peptides engineered to mimic the Tyr-His interactions of PSII [61], where very limited KSIEs accompany Tyr formation and oxidation [62]. PCET reactions can either proceed through step-wise or concerted mechanisms and have been shown to switch between these mechanisms depending on pH [57,63–66]. In the case of the Y191 FeCcP:Cc complex, potential vs proton-affinity dependencies suggest that the proton-coupled reaction likely proceeds with rate-limiting Y191 oxidation (ET) followed by sequential proton transfer (PT) [28]. However, in the ZnCcP system rates are only enhanced and radicals propagate to peripheral sites under conditions where a base aids the oxidation of the 191 relay site – a situation more likely to be governed by a concerted electron-proton transfer (CEPT) mechanism or sequential mechanism where PT contributes to the rate limiting step (**Figure 4**) [65,67]. Cationic Tyr radicals in proteins are extremely short lived (∼ 1 ps) and have low p*K*a values of ∼ -2 [68,69], thus once oxidized, PT from Y191^•+^ is unlikely to be rate limiting [70]. As discussed above, the relative orientations of the participating orbitals may prevent ZnP and W/Y191 from being well coupled, which would give rise to the switch it PCET mechanism in the photoexcited system, thereby demonstrating how small perturbations to the protein can have a substantial impact on multi-step ET reactions.

## Conclusions

Overall, our results demonstrate the detailed roles that PCET and hydrogen bonds play in modulating the reactivity of a hole-hopping relay site in a protein. Introduction of a proton acceptor for a tyrosyl radical rescues ET from Cc and alters radical propagation within ZnCcP. The proton acceptor has different functions depending on the relative potentials of the photogenerated acceptor (ZnP^•+^) and the ultimate donor (Cc or a peripheral protein site). The interacting conjugate base facilitates Tyr oxidation through PCET and tunes the resulting radical reduction potential through hydrogen bonding. In the case of peroxide-generated CpdI, which has high reactivity, Y191 oxidizes readily, but requires the hydrogen bond to retain a high enough potential to oxidize the donor site. In the case of the lower potential ZnP^•+^, the proton acceptor assists in deprotonation of the Tyr residue, effectively lowering its potential and thereby activating the relay site. An ensuing hydrogen bond to the oxidized Tyr^•^ again raises its potential to migrate the radical from the reaction center. These results also imply a reduction potential scale in the CcP system wherein CpdI > Y191^•+^ > ZnP^•+^ ∼ W191^•+^. For both FeCcP and ZnCcP, the high yield of W191^•+^ localizes the hole closer to the biological donor Fe(II) Cc, and is the primary electron acceptor on Cc. This work has implications for how proteins control the propagation of high-potential radical centers through hydrogen-bonding perturbations to relay sites, which, in principle, could be brought under allosteric regulation.

## Materials & Methods

### Protein expression and purification

CcP was expressed and purified as described previously [27,28]. *Escherichia coli* BL21(DE3) cells were transformed with the CcP gene in a ppSUMO vector and grown at 37°C in LB with 50 μg/mL kanamycin. When the OD600 nm reached between 0.7 and 0.9, cells were induced with 100 μM isopropyl β-D-1-thiogalactopyranoside (IPTG) and expressed at 20 °C overnight. Cells were lysed via sonication, and soluble protein was isolated by centrifugation. CcP was purified from lysate using a Ni-NTA column and His-SUMO tags were cleaved with ULP-1(R3) protease.

Tags were removed on a Ni-NTA column and CcP was collected in the flow through. Heme loading was low for the CcP variants as purified but could be fully realized after excess heme addition post-purification followed by additional isolation of the holo-protein. For FeCcP, the protein was stirred overnight at 4 °C with 1 molar equivalent of hemin dissolved in 0.1 M NaOH. The reaction was then neutralized with acetic acid and centrifuged to remove precipitated heme. For ZnCcP, the protein was stirred for 1 – 5 days at 4 °C with 2 – 5 molar excess of Zn protoporphyrin IX dissolved in DMF. Both FeCcP and ZnCcP were further purified via size exclusion chromatography and anion exchange chromatography. Fractions with A408nm/A280nm ≥ 1 for FeCcP and A432nm/A280nm ≥ 2 for ZnCcP representing the holo-protein were collected and concentrated. Proteins were then flash frozen in liquid nitrogen, and stored at -80 °C.

Cc was expressed in BL21(DE3) cells containing the cytochrome c maturation system, which is required for heme ligation and soluble holo-protein. Cell cultures were grown at at 37 °C in LB with 125 μg/mL ampicillin and 50 μg/mL δ-aminolevulenic acid, a precursor for heme production. The PBTR-1 vector with the Cc gene contains a *trc* promotor and autoinduces Cc expression during the growth. Cells were lysed via sonication and Cc was purified out of the lysate via cation exchange chromatography and size exclusion chromatography. Protein was concentrated, flash frozen, and stored at -80 °C.

### Global incorporation of 4-fluoro-glutamic acid

A tac promotor was cloned into the ppSUMO vector containing the CcP gene using primers tacF and tacR (**Table S1**). Overlap extension PCR cloning was performed and PCR products were digested with DpnI to remove the ppSUMO template. DNA sequences were confirmed via Sanger sequencing. F-CcP variants were expressed in *E. coli* PA340 cells from the Coli Genetic Stock Center collection which are auxotrophic for glutamic acid. Cells were grown at 37 °C in LB with 50 μg/mL kanamycin. When the OD600 reached ∼0.8 cells were spun down and washed with Nanopure water several times before resuspending in M9 minimal media with 50 μg/mL kanamycin and 100 μg/mL 4-fluoro-DL-glutamic acid (Key Organics, 95% purity). The cells were then induced with 100 μM isopropyl β-D-1-thiogalactopyranoside (IPTG) and expressed at 17 °C overnight. The F-Glu containing proteins were purified as described above, and the presence of F-Glu residues were confirmed via mass spectrometry.

### Multiple Turnover Reduction Kinetics

Cc was reduced with dithionite and buffer exchanged into a 100 mM KP*i* buffer at the appropriate pH. Multiple turnover measurements were conducted with 1800 μL of 1 μM CcP and 30 μM reduced Cc in 100 mM KP*i* buffer (pH 5 – 7) at 10 °C. An excess of Fe(II) Cc was used to minimize binding effects from Fe(III) Cc. Reactions were initiated by the rapid addition H2O2 while stirring at 700 RPM (Hewlett-Packard 89090A Peltier) with a final H2O2 concentration of 10 μM. The rate of Fe(II) Cc oxidation was monitored at the heme Q bands Abs550 nm - Abs540 nm (ε = 19.2 mM^-1^ cm^-1^) and, the CcP oxy-ferryl species at Abs434 nm (a Fe(II/III) Cc isosbestic point) using a Hewlett-Packard Agilent 8453 spectrophotometer. Given that peroxide was in excess and reaction between peroxide and CcP fast, at early times the reaction was considered under pseudo first-order conditions. Rate constants were obtained by fitting the Cc oxidation traces to monoexponential curves (MatLab; Mathworks):

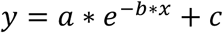

### Transient Absorption Spectroscopy (TA)

To avoid triplet state quenching from ground state oxygen, ZnCcP samples were prepared anaerobically. Cc was oxidized by potassium ferricyanide and was buffer exchanged into a degassed 10 or 100 mM KP*i* buffer (pH/D 6 – 8). For the KSIE studies, protein samples were left at 4 °C in their respective protonated or deuterated buffers for 1-2 hours to allow H/D exchange. Samples were prepared in a 1 mm cuvette with a 4 μL drop containing 75 μM ZnCcP and 75 – 150 μM Fe(III) Cc. ZnCcP alone was also measured for each variant to determine the rate of triplet state decay to the ground state (kD). Transient absorption measurements were performed as described previously [27]. Samples were placed in the path of a probe light, provided by a 75 W Xe-arc lamp and were excited at 560 nm by an Opotek Opolette Nd:YAG laser with approximately 2 mJ of power per 8 ns pulse firing at 20 Hz. The time delays between the excitation pump source and probe light were controlled by a Digital Delay Generator DG355 (Stanford Research Systems) which was triggered by the laser Q-switch. Time delays were varied between 0 to 40 ms randomly to reduce effects of photobleaching. Absorption from the probe light was measured using a Hamamatsu Photonic Multichannel Analyzer (PMA) and averaged 50 times. Difference spectra were calculated as Δ*A* = -log(excited/reference). Data were subjected to global analysis and singular value decomposition (SVD) was used to deconvolute the evolution-associated difference spectra (EADS) in Glotoran [71].

### Electron Paramagnetic Resonance Spectroscopy

Samples for Electron Paramagnetic Resonance (EPR) spectroscopy were prepared anaerobically in 100 mM KP*i* (pH 6 – 8) and 20 % glycerol buffer. A [Co(NH3)5Cl] Cl2 stock solution was prepared and degassed to act as a sacrificial electron acceptor for ^3^ZnP. 150 – 750 μM CcP was added to 150 – 1,500 μM [Co(NH3)5Cl] Cl2 for molar ratios ranging from 1:1 to 1:30. Around 20µL of the sample were inserted in Quartz capillary (Wilmad WG-221T-RB) with an internal diameter of 1.1 mm. The capillary were sealed with epoxy in a Coy Lab’s anaerobic chamber. Samples were either excited at 560 nm and flash-frozen or kept at room temperature and excited during collection for low and high temperature studies, respectively.

cw-EPR spectra were conducted using a Bruker E500 ElexSys spectrometer equipped with a ER 4123SHQE Bruker cavity. For 10 K measurements, the spectrometer was equipped with a liquid He cryostat ESR900 (Oxford Instrument). The acquisition parameters were fixed to 25 dB (0.63 mW) microwave power, 0.4 mT modulation amplitude and 100 kHz modulation frequency. For 110 K measurements, the temperature was stabilized by flushing cold nitrogen air through a transfer line to the cavity. The acquisition parameters were fixed to 40 dB (0.02 mW) microwave power, 0.5 mT modulation amplitude, 100 kHz modulation frequency, and a field sweep of 16.0 mT. For 298 K measurements, the acquisition parameters were 15 dB (6.35 mW) microwave power, 0.4 mT modulation amplitude, and 100 kHz modulation frequency. In situ irradiation was achieved with a 5 mW diode laser at λ = 532 ± 10 nm.

The power saturation curves were obtained at 110 K by varying the attenuation from 10 dB to 60 dB and recording each cw-EPR spectra individually. As the broad and the narrow components overlap, extraction of the peak-to-peak intensity was carried out by fitting each component with a Gaussian derivative using a global fitting procedure (see details in **Figure S2**). Half-saturation power P1/2 was obtained by fitting the peal to peak intensity Ipp(P) versus the microwave power with the following equation [72]:

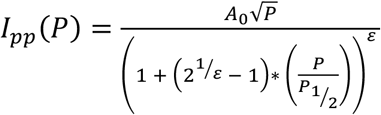

The ε parameter was fixed to 1.5 assuming a pure gaussian lineshape [72] and the amplitude A0 and the half-saturation power P1/2 were varied.

Pulse EPR experiments were conducted at 34 GHz using an Elexsys E580 spectrometer equipped with a 10 W solid state amplifier. All experiments were achieved in a Q-Band ENDOR EN 5107D2 Cavity. A Bruker E-580 AWG Arbitrary Waveform Generator was used for the microwave pulse generation. The temperature was varied by using an ER 4118HV-CF10-L FlexLine Cryogen-Free VT System. The pulse length (varying with the coupling range of the resonator and the microwave power used) was determined with a rabi nutation measurement π- τ2-π/2-τ-π-τ-echo by increasing sequentially the length of the first pulse.

The electron spin–lattice relaxation times T1 were measured over the temperature range by varying the repetition rate (or shot repetition time SRT) in two pulse echo sequence, π/2-τ- π-τ- echo. The magnetization recovery was fitted by systematically a mono (single radical species) and bi-exponential (double radical species) model. T1 values are reported in the **Table S8**. Phase- memory times Tm, were measured by a two-pulse echo decay sequence, π/2-τ-π-τ-echo, while varying the tau. The curves were fit by an exponential decay: I(t)=I0*exp(-2t/Tm) or biexponential decay I(t)=I1*exp(-2τ/Tm1) + I2*exp(-2τ/Tm2).

ENDOR (Electron Nuclear Double Resonance) measurements were conducted with a ^1^H Davies sequence π-t1-rf pulse-t2-π/2-τ-π-τ-echo as the hfccs of some protons can be very large for the tryptophanyl radical by instance (∼20 MHz for some positions). t1 and t2 were fixed to 1 µs to avoid the overlap of the rf pulse with microwave pulses. The rf pulse length has been fixed to 20 µs, length that shows the maximum proton NMR resonance intensity with the 150 W Bruker RF amplifier used. Selective π/2 and π pulses were used with a length around 70/140 ns respectively in accordance with a microwave power attenuation of 15 dB.

3-pulse ESEEM sequence π/2-τ-π/2-T-π/2-τ-echo was employed in Q-band to study the spin modulation of nitrogen couplings. To avoid any τ artifact, 12 slices of ESEEM was recorded with τ varying by steps of 10 ns and averaged during post-processing of the data. For every τ value, the 3P-ESEEM was recorded using a starting mixing time T = 100 ns and a dwell time of 8 ns to have sufficient frequency resolution in the area of 1-20 MHz. For this pulse experiment, the resonator was critically coupled and a length of 14 ns was employed for π/2 pulses.

Fitting of the cw-EPR spectra (**Figure S6-9**) were carried out by the EasySpin toolbox [73] under Matlab. The garlic.m function was employed to fit spectrum with 2 components. The first component with large linewidth was simulated with the use of four parameters: a fixed single ^1^H hyperfine coupling constant of 17 MHz, and three variables parameters: isotropic g-factor g1, a gaussian linewidth (full width half maximum) lw1, and a scaling factor A1. The second component with narrower linewidth was simulated with the use of three variables parameters: an isotropic g-factor g2, a gaussian linewidth (full width half maximum) lw2, and a scaling factor A2. The sum of the scaling factor was fixed to unity: A1 + A2 = 1.

The dipolar coupling line-broadening simulations (**Figure S4,5**) were carried out with the pepper.m function in EasySpin [73] with a first spin system of g1 = 2.0036, gaussian linewidth (full width half maximum) lw1 = 0.946 mT, an apparent ^1^H hyperfine coupling constant of 19 MHz and a second spin system g2 = 1.92, a gaussian linewidth (full width half maximum) lw2 = 0.946 mT. No hyperfine coupling was included for the second spin system and the g-factor was shifted to higher field to avoid overlap of the two components.

ESEEM and ENDOR spectra were simulated using saffron.m function of the EasySpin matlab toolbox [73].

### QM/MM Calculations and Analysis

The PDB file 1u74, 6p41, 5cih is used as the initial structure (provided in the github folder) for the W191, Y191:E232, and Y191 variants of ZnCcP:Cc. respectively. The protein forcefield is taken from CHARMM22 [74] and H molecules are added according to the topology file. The protein is solvated in a rectangular box of TIP3P water molecules. The size of the box is determined by the length of the protein; a 20 Å layer of water molecules is added around the protein (**Figure S15a-b**) and a 0.05 M buffer is added to make the system charge neutral. The solvent is then equilibrated with a 200-step steepest decent (SD) algorithm and a 1000-step Adopted Basis Newton-Raphson (ABNR) simulation. The structure obtained after MD equilibration are optimized at the QM/MM level using the software package Py-ChemShell [75], where QM calculations are performed with ORCA and the MD simulations use DL-POLY5.

Following previous experimental work and theoretical studies on the 2-step mechanism in the ZnCcP:Cc [27,39], we undertake QM/MM simulations to elucidate hole hopping between the ZnP and the W191/Y191 residues. The QM active site includes the porphyrin ring, axial His residue (H175), Asp (D235), Trp/Tyr (W/Y191, respectively), Leu/Glu (L/E232, respectively), and Trp (W51) as shown in **Figure S15c**. Initial QM/MM optimization with MM region frozen and using basis: 6-31G* with functionals B3LYP, WB97x and electrostatic embedding [76]. We find that the range-separated wB97x hybrid functional yields more accurate orbital energies and spin densities for the triplet and doublet states and choose this functional for the remaining calculations (**Figure S16** and accompanying text in the SI). We generate a 1ps long QM/MM-MD (NVT) trajectory for the W191, Y191, Y191:E232 variants with structures saved every 50 fs. Spin density analysis is performed using these snapshots to track hole localization (**Figures S17-S19**). To validate the pH-dependence studies, we run shorter (500 ps) trajectories for the protonated Y191:E232 and the deprotonated Y191 variants (**Figures S20-S21**).

### Supporting Information

The following files are available free of charge.

Additional details including DNA sequences and primers used for mutagenesis. Tables of fitted kinetic data for multiple-turnover and TA experiments. Tables for radical relaxation times and distributions in EPR measurements. XIC for MS of F-Glu containing peptides. Additional EPR data including controls, field swept echo for pulsed measurements, and simulations. Optimized MD structures, along with additional simulation details, motivation for choice of DFT functional, and QM/MM trajectory spin-density plots for all variants discussed in the manuscript. A GitHub Repository link is also provided where all relevant structure files, input files, and results of spin density analysis are included. (DOC)

## Author Information

## Authors

Rebecca K Zawistowski, Department of Chemistry and Chemical Biology, Weill Institute for Cell and Molecular Biology, Cornell University

Timothée Chauviré, Department of Chemistry and Chemical Biology, Weill Institute for Cell and Molecular Biology, National Biomedical Center for Advanced ESR Technologies (ACERT), Cornell University,

Sutanuka Manna, Department of Chemistry and Chemical Biology, Cornell University

Nandini Ananth, Department of Chemistry and Chemical Biology, Cornell University

## Acknowledgements

This work is supported by NSF grant MCB 2129729 to BRC and by ACERT under the NIH grant 1R24GM146107 and the NIH grant 1R35GM158400-01 to NA. We would like to thank Siddarth Chandrasekaran for the initial EPR measurements and analysis as well as Estella Yee for assistance with kinetics measurements.

