## Supplementary material for "Modulating radical propagation in proteins by proton-coupled electron transfer and hydrogen bonding": Table S1

### **Supporting Information for**: Modulating radical propagation in proteins by proton-coupled electron transfer and hydrogen bonding.

*Rebecca K. Zawistowski[1,2], Timothée Chauviré[1,2], Sutanuka Manna[1],*

*Nandini Ananth[1] and Brian R. Crane[1,2*]*

[1]Department of Chemistry and Chemical Biology,

[2]Weill Institute for Cell and Molecular Biology, Cornell University

Ithaca, NY 14853, USA

**Table of Contents:**


| | |
|---|---|
| 1-4. | DNA Sequence of tac-ppSUMO-CcP |
| 5. | Table S1. List of Primers |
| 6. | Table S2. Relative abundance of F-Glu232 |
| 7. | Table S3. Multiple turnover rate constants |
| 8-10. | Table S4-6. TA EADS rate constants |
| 11. | Table S7. Power saturation parameters |
| 12. | Table S8. Radical distribution at different $CoN_5$ ratios |
| 13. | Table S9. Radical relaxation times |
| 14 | Table S10 Simulation fitting parameters |
| 15 | Table S11. Radical distributions in W191 variants |
| 16. | Figure S1. MS data of 232-containing peptide |
| 17-18. | Figure S2-3. Power saturation analysis |
| 19-24. | Figure S4-9. cwEPR simulations |
| 25. | Figure S10. Pulsed EPR Spectra |
| 26. | Figure S11. Field swept echo for pulsed EPR spectra |
| 27. | Figure S12. $^1$H ENDOR simulations |
| 28-29. | Figure S13-14. $^{14}$N ESEEM simulations |
| 30. | Figure S15. Simulation Details: MD optimized structures and QM region |
| 31-32. | Figure S16. Choice of Functional |
| 33. | Figure S17. Spin Density along QM/MM trajectory for W191 |
| 34. | Figure S18. Spin Density along QM/MM trajectory for Y191:E232 |
| 35. | Figure S19. Spin Density along QM/MM trajectory for Y191 |
| 36. | Figure S20. Spin Density along QM/MM trajectory for Y191, Y191 deprotonated |
| 37. | Figure S21. Spin Density along QM/MM trajectory for Y191:E232, E232 protonated |
| 38-39. | Github Information |
| 40. | References |

**DNA sequence of tac-ppSUMO-CcP:**

TAATACGACTCACTATAGGTTGACAATTAATCATCGGCTCGTATAATGTGTGGAATT
GTGAGCGGATAACAATTCCCCTCTAGAAATAATTTTGTTTAACTTTAAGAAGGAGAT
ATACCATGGGCAGCAGCCATCATCATCATCATCACAGCAGCGGCCTGGTGCCGCGC
GGCAGCCATATGGCTAGCATGTCGGACTCAGAAGTCAATCAAGAAGCTAAGCCAGA
GGTCAAGCCAGAAGTCAAGCCTGAGACTCACATCAATTTAAAGGTGTCCGATGGAT
CTTCAGAGATCTTCTTCAAGATCAAAAAGACCACTCCTTTAAGAAGGCTGATGGAAG
CGTTCGCTAAAAGACAGGGTAAGGAAATGGACTCCTTAAGATTCTTGTACGACGGT
ATTAGAATTCAAGCTGATCAGACCCCTGAAGATTTGGACATGGAGGATAACGATATT
ATTGAGGCTCACAGAGAACAGATTGGTGGATCCATGATCACCACGCCGCTCGTTCAT
GTCGCCTCTGTCGAAAAAGGGAGGTCATACGAGGACTTCCAAAAGGTGTACAATGC
GATTGCACTCAAGCTGAGGGAAGATGACGAATATGACAACTATATAGGCTATGGGC
CCGTATTAGTCCGTCTTGCTTGGCACATTTCAGGGACCTGGGACAAGCACGACAATA
CAGGCGGGTCATACGGTGGTACATACAGATTCAAAAAGGAGTTTAACGATCCATCC
AATGCGGGCTTGCAGAATGGCTTTAAGTTCCTGGAGCCCATTCACAAAGAGTTTCCC
TGGATCTCCTCGGGTGATCTGTTCAGTCTAGGGGGTGTCACTGCTGTGCAGGAAATG
CAGGGTCCCAAGATTCCATGGAGATGTGGTAGAGTCGACACGCCAGAGGATACTAC
CCCTGACAACGGGAGACTGCCTGACGCTGATAAGGACGCTGGCTATGTCAGAACGT
TTTTTCAAAGACTTAATATGAATGACAGAGAAGTAGTTGCTCTTATGGGGGCTCACG
CTCTGGGCAAGACCCACTTGAAGAACTCTGGATACGAAGGGCCATACGGAGCCGCT
AACAACGTCTTTACCAATGAGTTTTACTTGAACTTGTTGAATGAAGACTGGAAATTG
GAAAAGAACGACGCGAACAACGAACAGTGGGACTCTAAGAGCGGCTACATGATGCT
GCCCACTGATTATTCTTTGATTCAAGATCCCAAGTACTTAAGCATTGTGAAAGAATA

CGCTAATGACCAGGACAAGTTCTTCAAGGATTTTTCCAAAGCTTTTGAAAAACTGTT
GGAAAACGGTATCACTTTCCCTAAAGACGCGCCCAGTCCATTTATTTTCAAGACTTT
AGAGGAACAAGGTTTATAG

| Name | Sequence 5' – 3' |
|---|---|
| tacF | CCCGCGAAATTAATACGACTCACTATAGGTTGACAATTAATCATCGGCTCG TATA<br><br>ATGT |
| tacR | GGGGAATTGTTATCCGCTCACAATTCCACACATTATACGAGCCGATGATTA ATTG |

**Table S1.** List of Primers

| **Peptide** | **Peak area ($x10^9$)** | **%** |
|---|---|---|
| F-Glu-peptide 2+ | 2.18 | 48.34 |
| Glu peptide 2+ | 2.33 | 51.66 |
| Total | 4.52 | 100.00 |

**Table S2.** Relative abundance of F-Glu 232 containing peptide prior to heme-conjugation

| **Variant** | **pH** | ***A*** | $\boldsymbol{k_1}$ **[s$^{-1}$]** |
|---|---|---|---|
| Y191 CcP | 5 | 0.496 ± 0.047 | 0.0055 ± 0.0009 |
| | 6 | 0.428 ± 0.051 | 0.0043 ± 0.0014 |
| | 7 | 0.437 ± 0.014 | 0.0060 ± 0.0004 |
| Y191:E232 CcP | 5 | 0.573 ± 0.038 | 0.0649 ± 0.018 |
| | 6 | 0.525 ± 0.040 | 0.0571 ± 0.002 |
| | 7 | 0.526 ± 0.018 | 0.0294 ± 0.009 |
| Y191 F-$E_G$ CcP | 5 | 0.399 ± 0.104 | 0.0095 ± 0.0032 |
| | 6 | 0.281 ± 0.016 | 0.011 ± 0.001 |
| | 7 | 0.415 ± 0.010 | 0.0041 ± 0.001 |
| Y191:F-E232 F-$E_G$ CcP | 5 | 0.474 ± 0.070 | 0.0458 ± 0.0093 |
| | 6 | 0.468 ± 0.116 | 0.0292 ± 0.0055 |
| | 7 | 0.455 ± 0.014 | 0.0114 ± 0.0014 |
| 85% Y191:E232 CcP + 15% Y191 CcP | 6 | 0.546 ± 0.015 | 0.0386 ± 0.0071 |
| | 7 | 0.452 ± 0.046 | 0.0297 ± 0.0045 |

**Table S3.** Multiple turnover rate constants in 100 mM $KP_i$ buffer. Variations in the scaler coefficients, A, may vary somewhat due to fluctuations in the initial protein concentrations.

| Variant | pH | $k_p$ [$s^{-1}$] | $k_e$ [$s^{-1}$] | $k_{eb}$ [$s^{-1}$] | $k_D$ [$s^{-1}$] |
|---|---|---|---|---|---|
| Y191 CcP:Cc | 6 | 319 ± 10 | 219 ± 12 | 25 ± 3 | 100 ± 6 |
| | 7 | 296 ± 26 | 188 ± 27 | 21 ± 4 | 107 ± 8 |
| | 7.5 | 263 ± 11 | 176 ± 13 | 36 ± 3 | 87 ± 7 |
| | 8 | 287 ± 21 | 191 ± 22 | 50 ± 17 | 96 ± 7 |
| Y191:E232 CcP:Cc | 6 | 562 ± 57 | 399 ± 57 | 17 ± 2 | 163.0 ± 1.3 |
| | 7 | 509 ± 52 | 402 ± 53 | 79 ± 11 | 107 ± 6 |
| | 7.5 | 492 ± 18 | 400 ± 18 | 98.2 ± 0.5 | 92 ± 4 |
| | 8 | 285 ± 64 | 199 ± 64 | 104 ± 18 | 87 ± 8 |
| Y191:H232 CcP:Cc | 6 | 318 ± 22 | 207 ± 22 | 29 ± 3 | 112 ± 3 |
| | 7 | 367 ± 33 | 251 ± 33 | 34 ± 4 | 116 ± 4 |
| | 7.3 | 307 ± 36 | 212 ± 36 | 65 ± 5 | 95 ± 4 |
| | 7.5 | 294 ± 32 | 190 ± 32 | 73 ± 2 | 104 ± 8 |
| | 8 | 363 ± 34 | 260 ± 34 | 87 ± 10 | 103 ± 4 |

**Table S4.** Fitted rate constants from TA EADS in 100 mM $Kp_i$ buffer. $k_e$ is calculated from the difference between $k_p$ and $k_D$.

| Variant | pH | $k_p$ [$s^{-1}$] | $k_e$ [$s^{-1}$] | $k_{eb}$ [$s^{-1}$] | $k_D$ [$s^{-1}$] |
|---|---|---|---|---|---|
| Y191 CcP:Cc | 6 | 321 ± 23 | 220 ± 23 | 29 ± 2 | 101 ± 3 |
| | 7 | 328 ± 33 | 216 ± 34 | 37 ± 3 | 112 ± 8 |
| | 8 | 389 ± 39 | 294 ± 40 | 68 ± 5 | 95 ± 7 |
| Y191:E232 CcP:Cc | 6 | 325 ± 15 | 306 ± 16 | 19 ± 2 | 149 ± 6 |
| | 7 | 447 ± 16 | 320 ± 16 | 127 ± 23 | 109 ± 1 |
| | 8 | 606 ± 99 | 406 ± 99 | 200 ± 27 | 84 ± 4 |

**Table S5.** Fitted rate constants from TA EADS in 10 mM $Kp_i$ buffer. $k_e$ is calculated from the difference between $k_p$ and $k_D$.

| Variant | pH | $k_p$ [$s^{-1}$] | $k_e$ [$s^{-1}$] | $k_{eb}$ [$s^{-1}$] | $k_D$ [$s^{-1}$] | $k_{H^+}/k_{D^+}$ |
|---|---|---|---|---|---|---|
| Y191 CcP:Cc | 7 | 296 ± 26 | 188 ± 27 | 21 ± 4 | 107 ± 8 | 1.05 |
| | 7 $D_2O$ | 348 ± 66 | 265 ± 67 | 20 ± 3 | 83 ± 8 | |
| Y191:E232 CcP:Cc | 6 | 561 ± 30 | 373 ± 31 | 24 ± 4 | 143 ± 6 | 1.04 |
| | 6 $D_2O$ | 399 ± 13 | 309 ± 14 | 23 ± 2 | 90 ± 4 | |
| | 7 | 631 ± 23 | 522 ± 23 | 122 ± 8 | 109 ± 3 | 1.71 |
| | 7 $D_2O$ | 582 ± 13 | 498 ± 23 | 71.2 ± 1.8 | 83 ± 19 | |
| Y191:H232 CcP:Cc | 7 | 370 ± 37 | 264 ± 37 | 37 ± 2 | 106 ± 3 | 0.949 |
| | 7 $D_2O$ | 295 ± 22 | 220 ± 27 | 39 ± 2 | 75 ± 15 | |
| | 7.3 | 307 ± 36 | 212 ± 36 | 65 ± 5 | 95 ± 4 | 1.77 |
| | 7.3 $D_2O$ | 309 ± 11 | 246 ± 11 | 37 ± 8 | 63 ± 2 | |
| | 7.5 | 274 ± 9 | 170 ± 9 | 69 ± 8 | 104 ± 3 | 1.50 |
| | 7.5 $D_2O$ | 223 ± 39 | 145 ± 39 | 46 ± 20 | 78 ± 2 | |

**Table S6.** Fitted rate constants from TA EADS. $k_e$ is calculated from the difference between $k_p$ and $k_D$. $k_{H^+}/k_{D^+}$ is the calculated ratio between $k_{eb}$ values in protonated vs. deuterated buffers.

| **Component** | $\mathbf{A_0}$ | $\mathbf{P_{1/2}}$ **(mW)** | **ε** | $\mathbf{R_2}$ |
|---|---|---|---|---|
| Broad ($W^{\bullet +}$) | 50 ± 14 | 0.27 ± 0.15 | 1.5 | 0.632 |
| Narrow ($ZnP^{\bullet +}$) | 177 ± 3 | 10.30 ± 0.33 | 1.5 | 0.999 |

**Table S7.** Fitting parameters for power saturation studies in **Figures S2,3**.

| Ratio CcP:$CoN_5$ | $X^{\bullet}$ | | | $ZnP^{\bullet+}$ | | |
|---|---|---|---|---|---|---|
| | **(%)** | **Linewidth (mT)** | **g-factor*** | **(%)** | **Linewidth (mT)** | **g-factor*** |
| 1:1 | 93.5 ± 0.1 | 1.18 ± 0.02 | 2.0034 ± 0.0002 | 6.5 ± 0.8 | 0.56 ± 0.03 | 2.0018 ± 0.0004 |
| 1:2 | 88.4 ± 0.1 | 1.31 ± 0.02 | 2.0033 ± 0.0002 | 11.6 ± 0.4 | 0.58 ± 0.01 | 2.0024 ± 0.0002 |
| 1:10 | 76.5 ± 0.1 | 1.63 ± 0.02 | 2.0026 ± 0.0004 | 23.5 ± 0.4 | 0.576 ± 0.003 | 2.0026 ± 0.0002 |
| ZnP | - | - | - | - | 0.49 ± 0.02 | 2.0023 ± 0.0002 |

**Table S8.** Radical distributions and linewidth (mT) in W191 ZnCcP samples at 110 K post-light excitation. Uncertainty corresponds to ± 2 σ and were obtained with the esfit.m function in easyspin. *g-factors were extracted by using the center field obtained by the two components fit following the procedure described by Bolton et al [1]. The 2,2-diphenyl-1-picrylhydrazyl (DPPH) was used as a g-factor standard ($g_{DPPH}$ = 2.0036 ± 0.0001).

| Variant | Temperature (K) | pH | $T_1$ | | $T_m$ | |
|---|---|---|---|---|---|---|
| | | | Trp/X | ZnP | Trp/X | ZnP |
| W191 | 20 | 6 | > 35 ms | 1.5 to 16 ms | 2.7 -3.0 μs | 1.6 – 2.2 μs |
| | 110 | 6 | > 5 ms | 80-500 μs | 2.5 -2.8 μs | 0.8 – 2.5 μs |
| G191 | 60 | 6 | na | 489 μs | na | 1.32 μs |
| | 110 | 6 | na | 206 μs | na | 797 ns |
| Y191:E232 | 80 | 7 | 3.2 ms | 425 μs | 2.0 ms | 381 μs |
| | 60 | 8 | 6.1 ms | 206 μs | 1.8 - 2.0 ms | 302-443 μs |
| | 110 | 7 | 2.3 ms | 301 μs | 1.9 ms | 560 μs |
| | 110 | 8 | 2.6 ms | 421 μs | 2.1 ms | 556 μs |

**Table S9.** Radical relaxation times in ZnCcP variants

| Sample | Component | Linewidth (mT) | (%) |
|---|---|---|---|
| With irradiation | $Trp^{\bullet+}$ | 1.15 ± 0.02 | 72.8 ± 0.3 |
| | $ZnP^{\bullet+}$ | 0.57 ± 0.004 | 27.2 ± 0.7 |
| Post-irradiation | $Trp^{\bullet+}$ | 0.89 ± 0.01 | 100 ± 0.0 |
| | $ZnP^{\bullet+}$ | - | - |

**Table S10.** Simulation parameters for Figure S4

| Variant | pH | X$^{\bullet}$ | | ZnP$^{\bullet+}$ | |
|---|---|---|---|---|---|
| | | (%) | Linewidth (mT) | (%) | Linewidth (mT) |
| W191 | 6 | 0.743 ± 0.003 | 1.20 ± 0.02 | 0.257 ± 0.008 | 0.57 ± 0.01 |
| | 7 | 0.485 ± 0.006 | 1.13 ± 0.02 | 0.515 ± 0.006 | 0.57 ± 0.01 |
| | 8 | 0.688 ± 0.006 | 1.10 ± 0.03 | 0.312 ± 0.012 | 0.56 ± 0.01 |
| Y191 | 6 | 0.473 ± 0.015 | 0.97 ± 0.04 | 0.527 ± 0.013 | 0.55 ± 0.01 |
| | 7 | 0.560 ± 0.006 | 1.08 ± 0.03 | 0.440 ± 0.008 | 0.56 ± 0.01 |
| | 8 | 0.848 ± 0.002 | 1.04 ± 0.03 | 0.152 ± 0.008 | 0.54 ± 0.01 |
| Y191:E232 | 6 | 0.613 ± 0.006 | 1.14 ± 0.03 | 0.387 ± 0.010 | 0.58 ± 0.01 |
| | 7 | 0.929 ± 0.001 | 1.06 ± 0.02 | 0.071 ± 0.008 | 0.52 ± 0.03 |
| | 8 | 0.967 ± 0.001 | 1.14 ±0.03 | 0.033 ± 0.006 | 0.51 ± 0.05 |
| Y191:H232 | 6 | 0.958 ± 0.001 | 1.03 ± 0.02 | 0.042 ± 0.006 | 0.53 ± 0.04 |
| | 7 | 0.944 ± 0.001 | 1.03 ± 0.02 | 0.056 ± 0.007 | 0.54 ± 0.04 |
| | 8 | 0.982 ± 0.001 | 1.01 ± 0.02 | 0.018 ± 0.004 | 0.50 ± 0.06 |

**Table S11.** Radical distributions and linewidth (mT) in ZnCcP samples at 293 K post-light excitation. Uncertainty corresponds to ± 2 σ and were obtained with the esfit.m function in easyspin.

**a)** **ANNEQWDSKSGYMM(fluoro)EPTDYSLIQDPKYLS**

**b)**

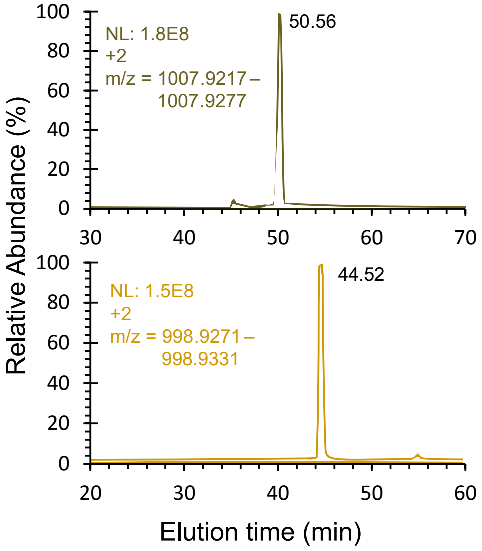


**Figure S1. a)** AspN digestion cleavage sites for E232 containing peptide. **b)** Extracted ion chromatogram of peptide DSKSGYMMEPTDYSLIQ. The F-Glu containing peptide (Top) elutes later than the native Glu residue (bottom).

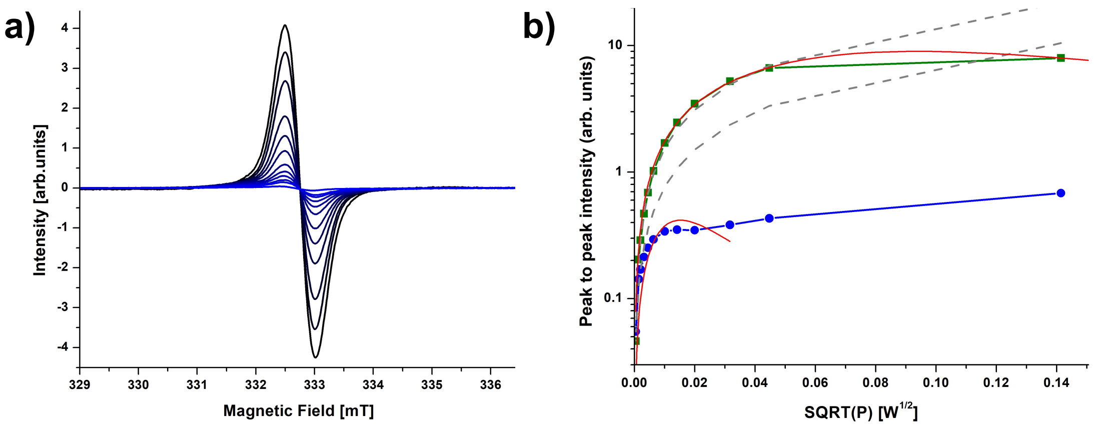


**Figure S2.** cwEPR power saturation analysis conducted at 110 K of W191 ZnCcP at pH6. **a)** cwEPR spectra recorded at different power attenuation varying from 10 dB (0.02 W) (darkest blue) to 60 dB ($2x10^{-7}$ W) (lightest blue). **b)** Peak to peak intensity (Ipp) of the extracted narrow (green) and broad components (blue) versus the square root (SQRT) of the microwave power. Gray dashes curve represent the peak-to-peak intensity varying linearly with the square root of P in the case of a non-saturating radical. Red curves are the fits obtained with the following equation:

$$I_{pp}(P) = \frac{A_0\sqrt{P}}{\left(1 + (2^{1/\varepsilon} - 1) * \left(\frac{P}{P_{1/2}}\right)\right)^{\varepsilon}}$$

For the narrow component, the fit was carried out for the whole set of datapoints. The ε parameter was fixed to 1.5 assuming a pure gaussian line shape [1]. For the broad component, the 2 last points were excluded of the fit assuming the intensity of the broad component were poised by the narrow component. Half-saturation power $P_{1/2} = 0.27 \pm 0.15$ mW (slow relaxing radical) and $P_{1/2} = 10.30 \pm 0.33$ mW (fast relaxing radical) were found respectively for the broad and the narrow components indicating the presence of distinct radicals with different microwave saturation behavior.

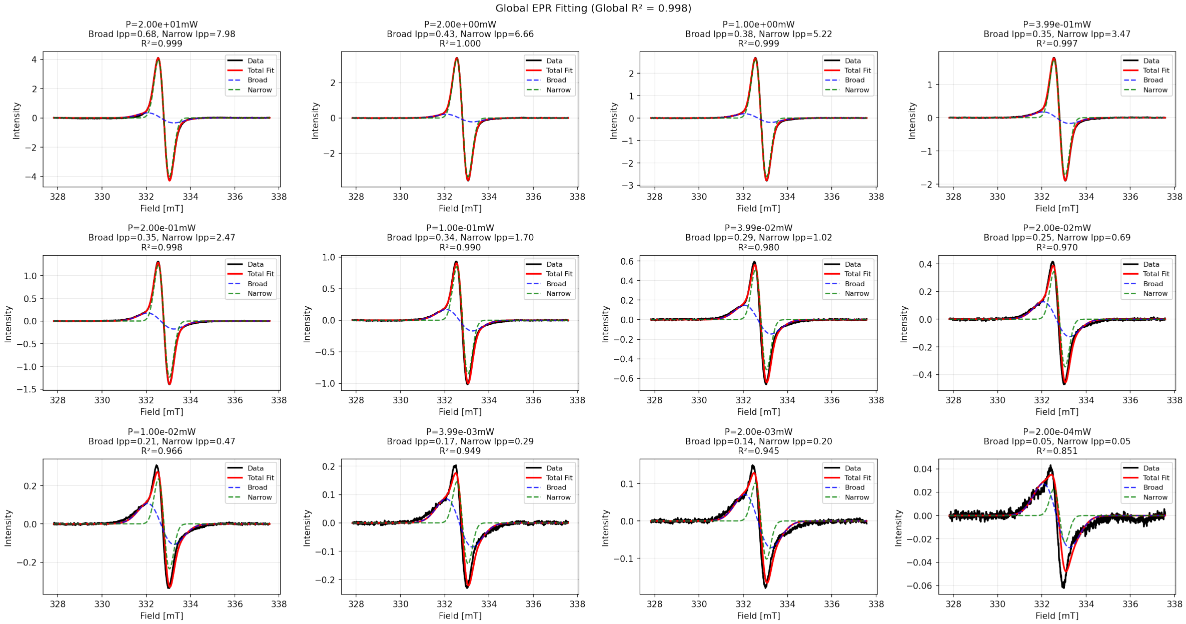


**Figure S3.** Global fitting procedure to extract the peak-to-peak intensity of the broad (blue dash line) and narrow component (green dash line) for power saturation analysis in **Figure S2**. The sum of the fitted component is plotted as the red line and compare to the black experimental spectra. For each component, the fit was carried out using a Gaussian derivative function:

$$I(B) = -1 * e^{\frac{1}{2}} * A_0 * \frac{B - B_{center}}{0.5 * lwpp} * e^{-1/2\left(\frac{B - B_{center}}{0.5 * lwpp}\right)^2}$$

where B is the microwave field, $A_0$ is an amplitude factor, $B_{center}$ is the center field, and lwpp is the linewidth peak-to-peak. For each component, the center field and linewidth peak-to-peak were fixed to: 332.6566 mT and 1.196 mT respectively for the broad component and 332.8003 mT and 0.504 mT respectively for the narrow component.

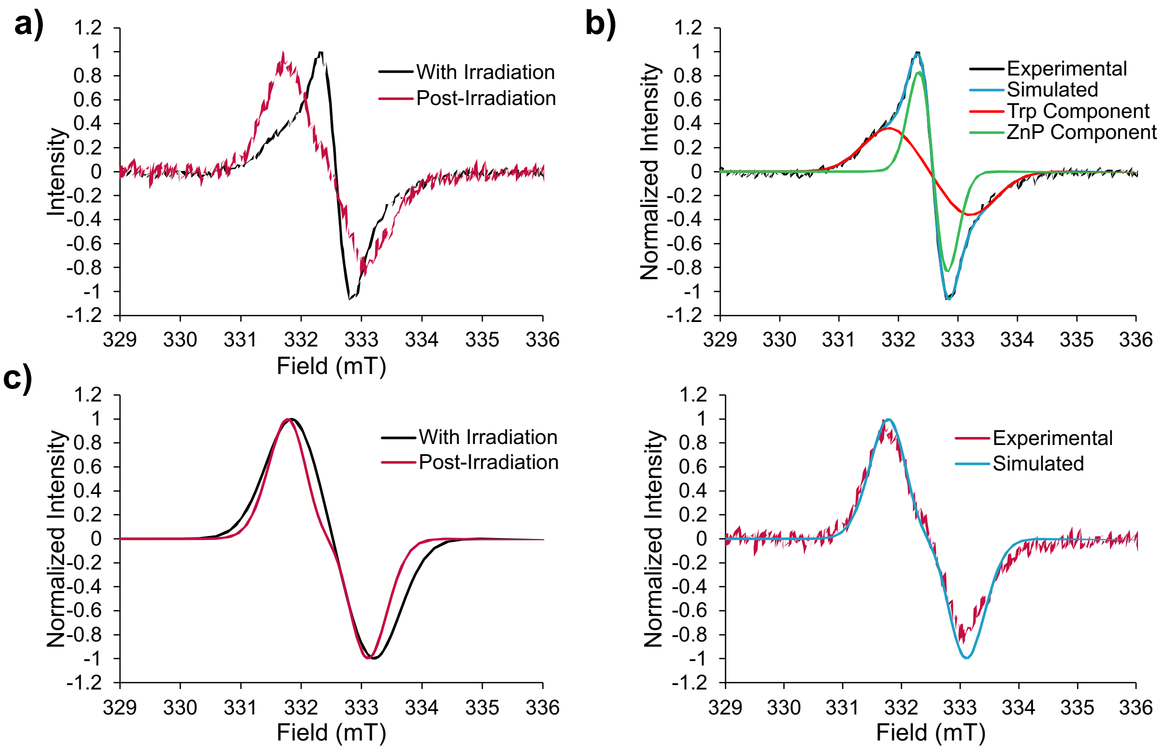


**Figure S4.** Evolution of the cwEPR spectra of W191 ZnCcP with stoichiometric amounts of $CoN_5$ at 293 K with and without irradiation (λ = 532 nm). **a)** Experimental cwEPR spectra during irradiation (black) and stored in dark 1 hr post-irradiation (red). **b)** Double component fits of the normalized data recorded during irradiation (top). Single component fits of the normalized data recorded post-irradiation (bottom). Double component fits were attempted and unsuccessful, indicating the presence of a single radical species only. Simulations were carried out by using the function garlic.m in easyspin. See Table S10 for simulation parameters. An additional apparent isotropic $^{1}H$ hyperfine coupling of 1.9 mT was used to reproduce the shape of the broad component. **c)** Normalized comparison of the broad components obtained from fitting in panel b) during irradiation (black) and after irradiation (red). A slight broadening of the broad component seems to be observed with the presence of the narrow component as indicated by an increase of 0.26 mT of the linewidth broadening (see Table S10).

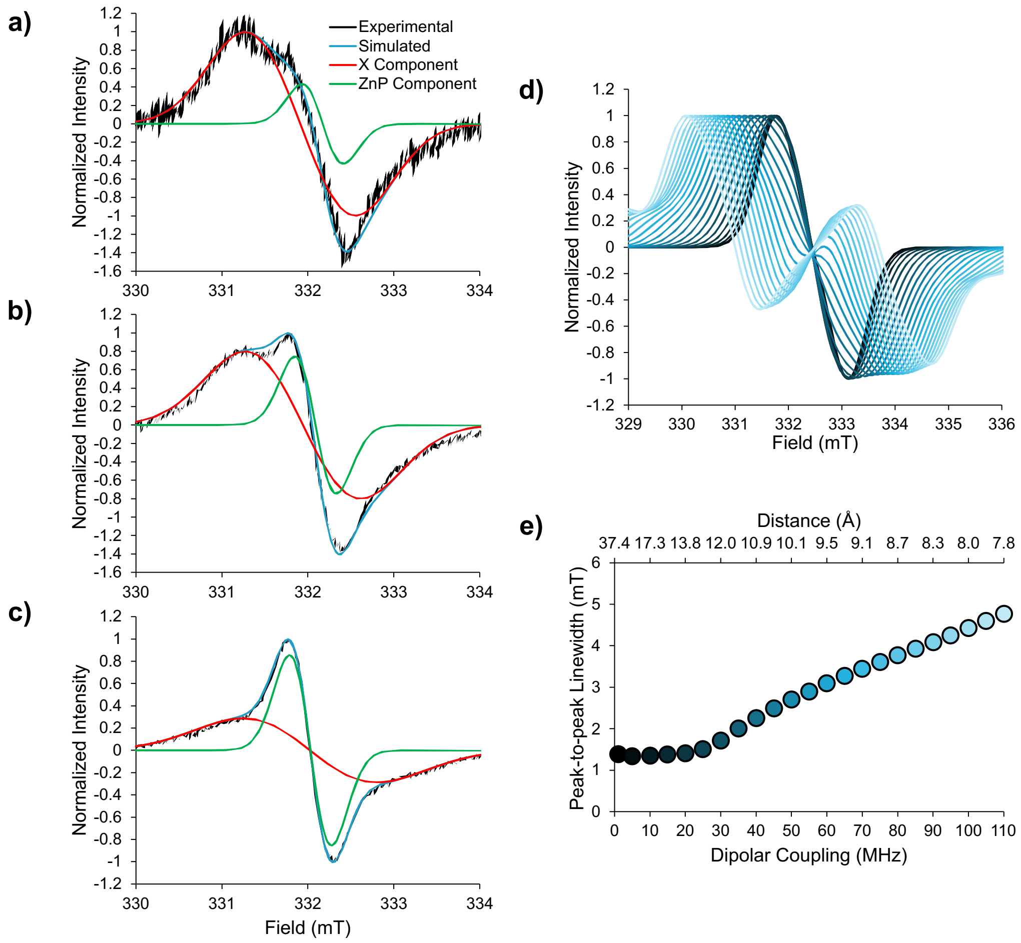


**Figure S5.** Experimental and simulated cwEPR spectra of **a)** 1:1 **b)** 1:2 and **c)** 1:10 W191 ZnCcP:$CoN_5$ at 110 K. **d)** Simulated cw-EPR spectra of two radicals with an electron spin S=1/2 interacting by dipolar coupling. The dipolar coupling was varied from 1 (black) to 110 MHz (blue) by steps of 5 MHz. Spectra were normalized by the maximum value of the first peak. A dipolar coupling of 110 MHz, representative of two spins separated by 7.8 Å (distance calculated from the point dipole approximation) would induce a peak-to-peak linewidth in the range of 4-5 mT. **e)** Extracted peak-to peak-linewidth from simulations in figure d). Distances were calculated using the point dipole approximation formula: $r = \left(\frac{\mu_0 g_1 g_2 \mu_B^2}{4\pi h \nu_{DD}}\right)^{1/3}$ where $\mu_0$ is the vacuum permeability, $\mu_B$ is the Bohr magneton, $g_1$ and $g_2$ are the g-factors of the two electron spins, h is Planck's constant, and $\nu_{DD}$ is the frequency of the dipolar coupling. See Methods in main text for simulation parameters.

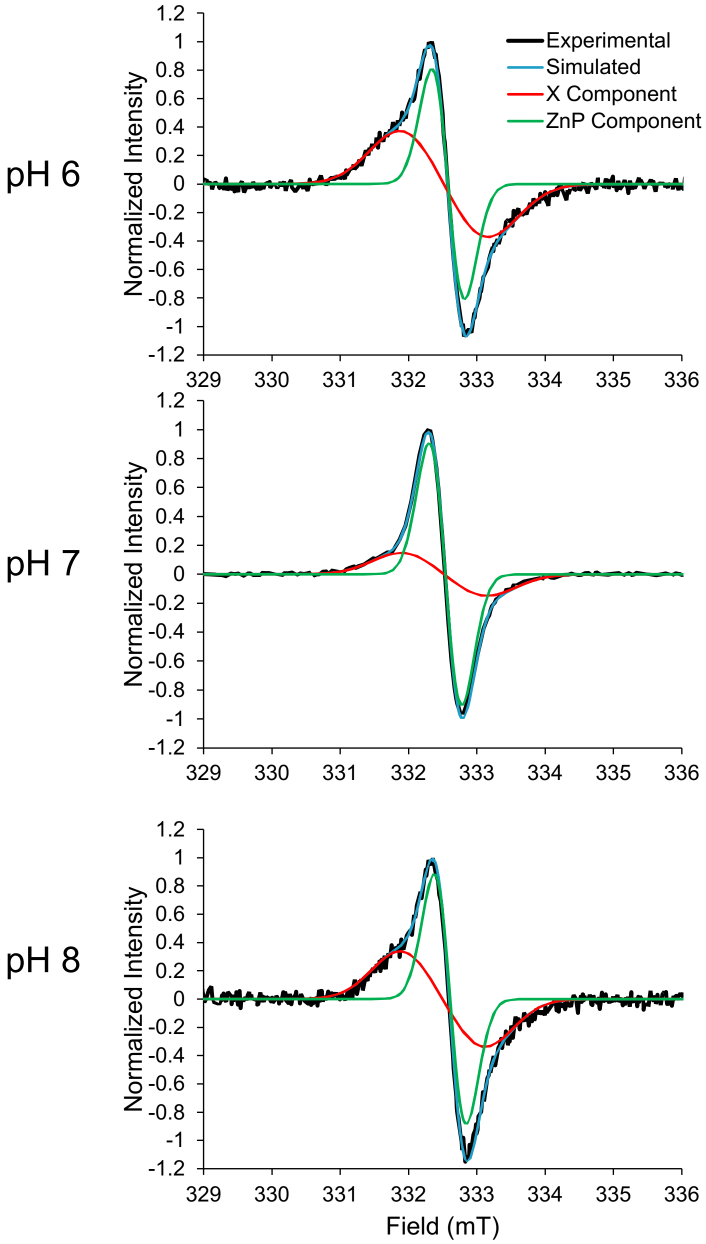


**Figure S6.** Simulated cwEPR spectra of W191 ZnCcP with $CoN_5$ at 293 K. See Methods for simulation parameters.

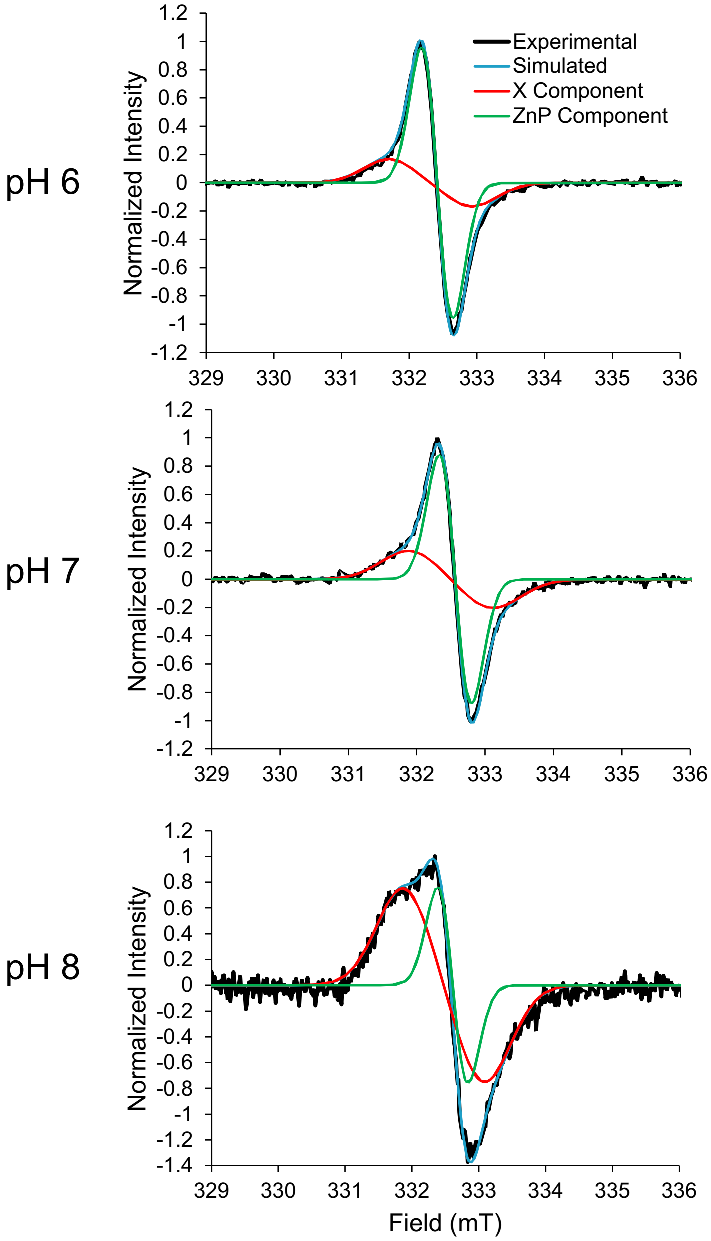


**Figure S7.** Simulated cwEPR spectra of Y191 ZnCcP with $CoN_5$ at 293 K. See Methods for simulation parameters.

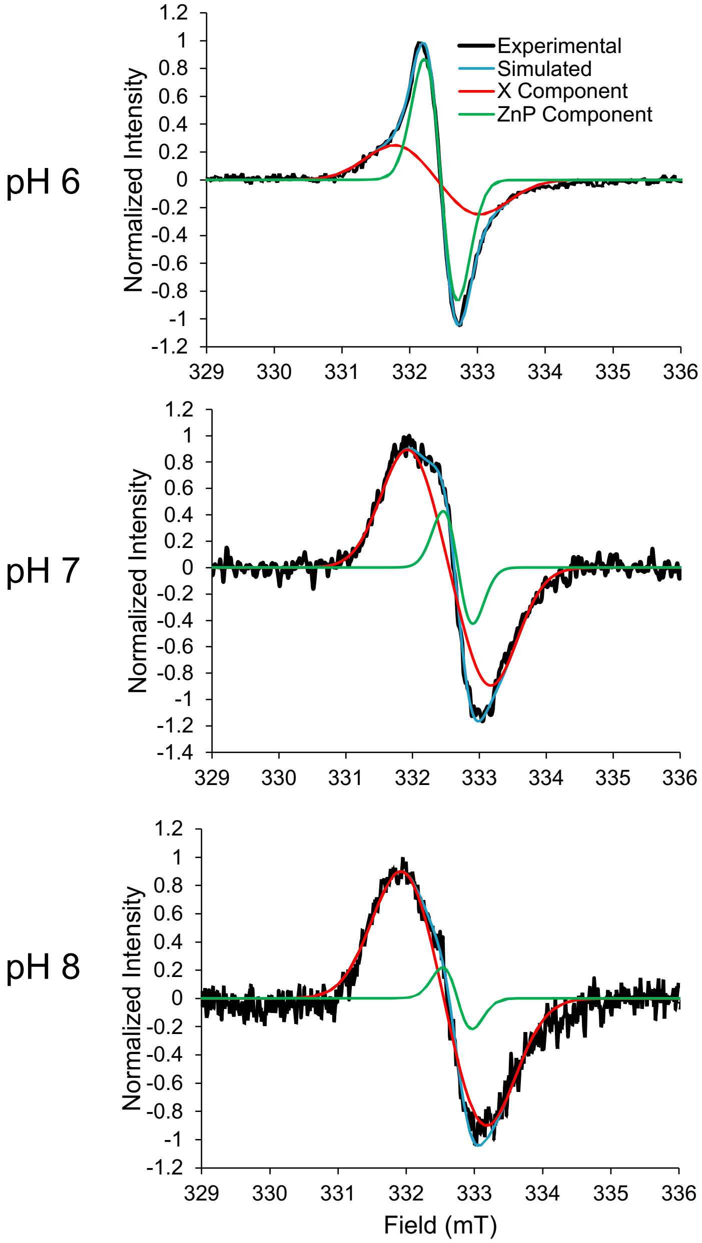


**Figure S8.** Simulated cwEPR spectra of Y191:E232 ZnCcP with $CoN_5$ at 293 K. See Methods for simulation parameters.

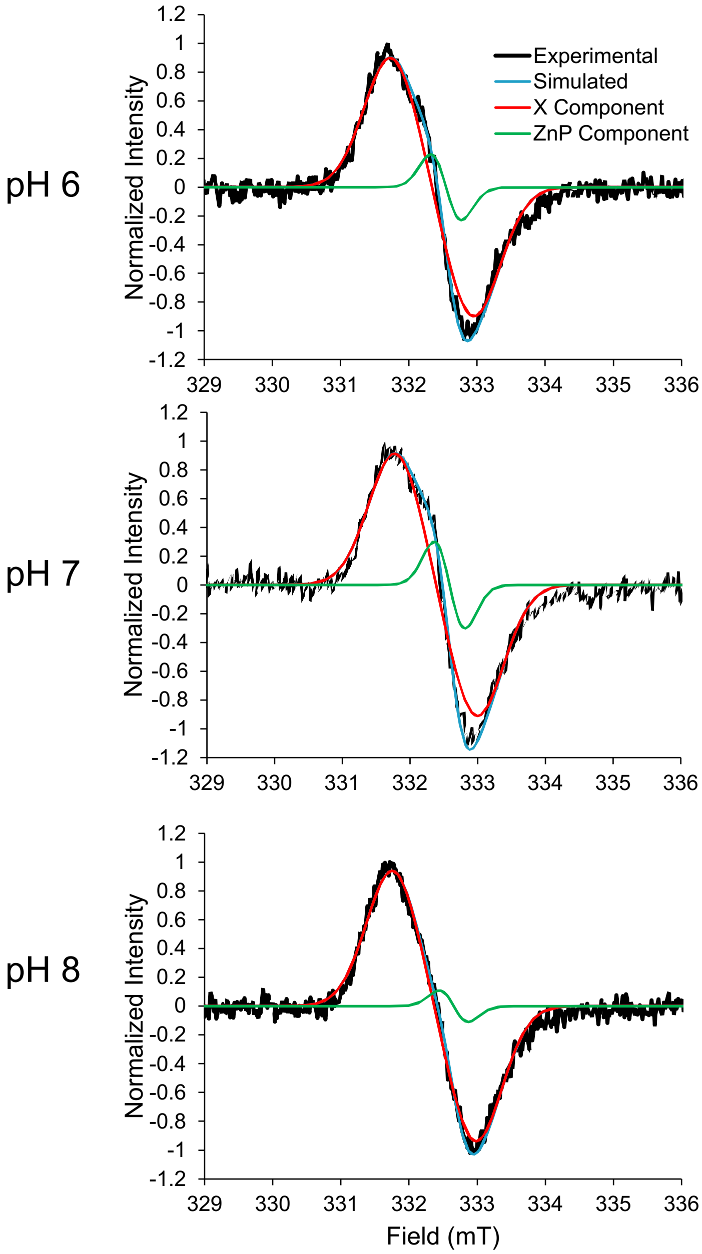


**Figure S9.** Simulated cwEPR spectra of Y191:H232 ZnCcP with $CoN_5$ at 293 K. See Methods for simulation parameters.

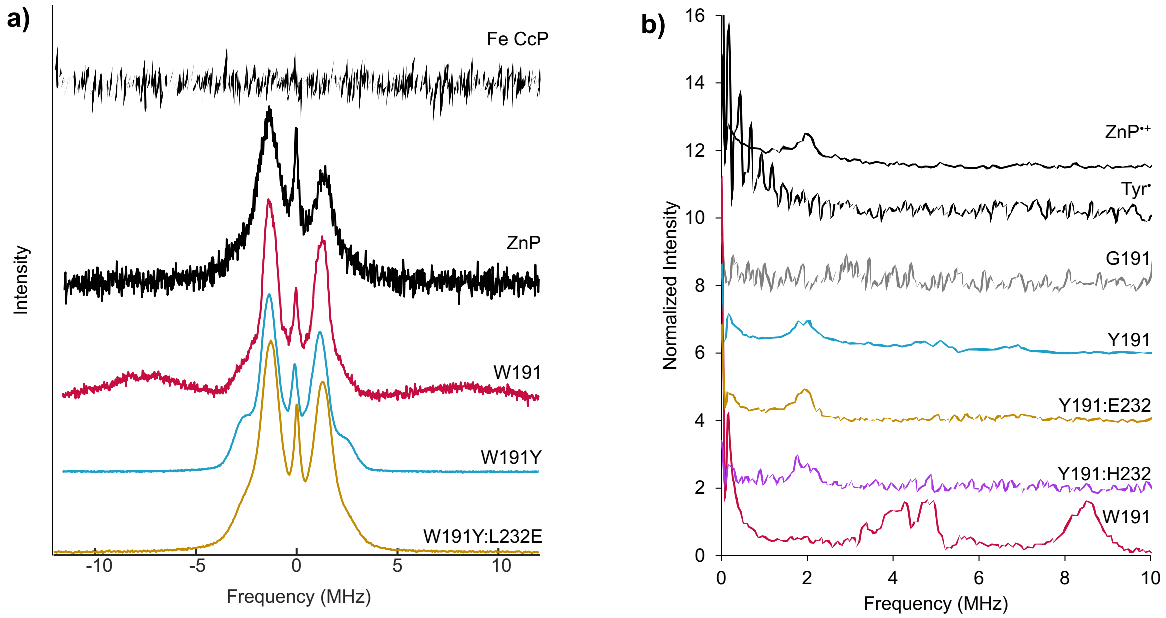


**Figure S10.** Hyperfine spectroscopy of CcP radical species. **a)** $^{1}H$ ENDOR spectra of FeCcP, free $ZnP^{\bullet+}$, ZnCcP variants at 10 K, pH 6 with 10-fold excess $CoN_5$. Central peaks representing the proton Larmor frequency are present in spectra of all ZnCcP species. Signals from hyperfine-shifted $Trp^{\bullet+}$ protons are also present in W191 ZnCcP at 15 MHz. wt FeCcP is absent of any peaks in the spectra indicating that $ZnP^{\bullet+}$ generation is necessary for oxidizing Trp191. **b)** $^{14}N$ ESEEM spectra at 110 K, pH 6 with 10-fold excess $CoN_5$ centered on the low-field radical transition. A sharp peak at 2 MHz is attributed to hyperfine interactions with the porphyrin nitrogen. 5 and 7-8 MHz peaks from $^{14}N$-coupling in Trp is less prominent at this temperature for the Y191 variants.

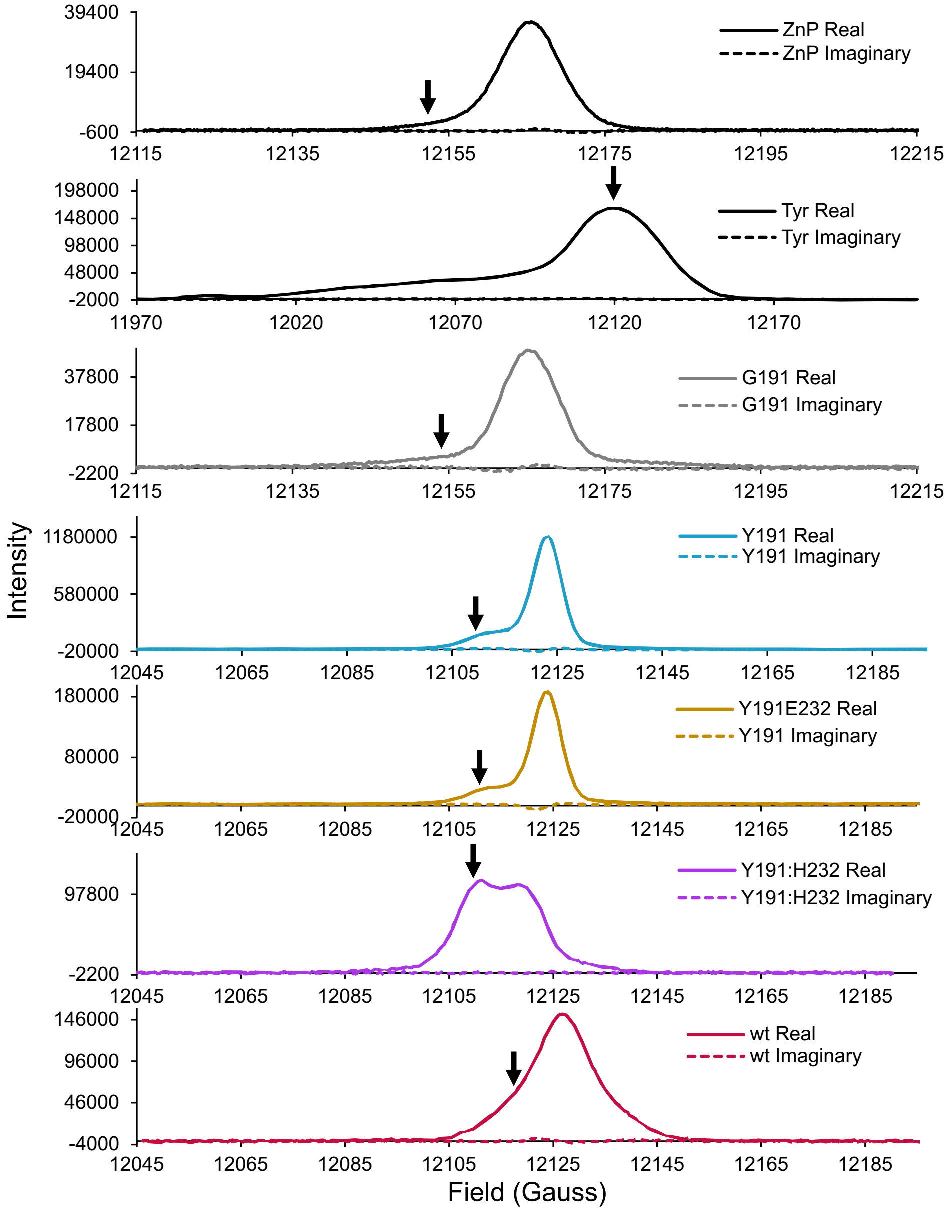


**Figure S11.** Field Swept Echos (FSE) of $^{1}$H ENDOR and $^{14}$N ESEEM spectra. Arrow indicates field position at which the EPR spectra (see **Figure 7** in main text) were acquired. The magnetic field shift observed between spectra is induced by the experimental frequency shift.

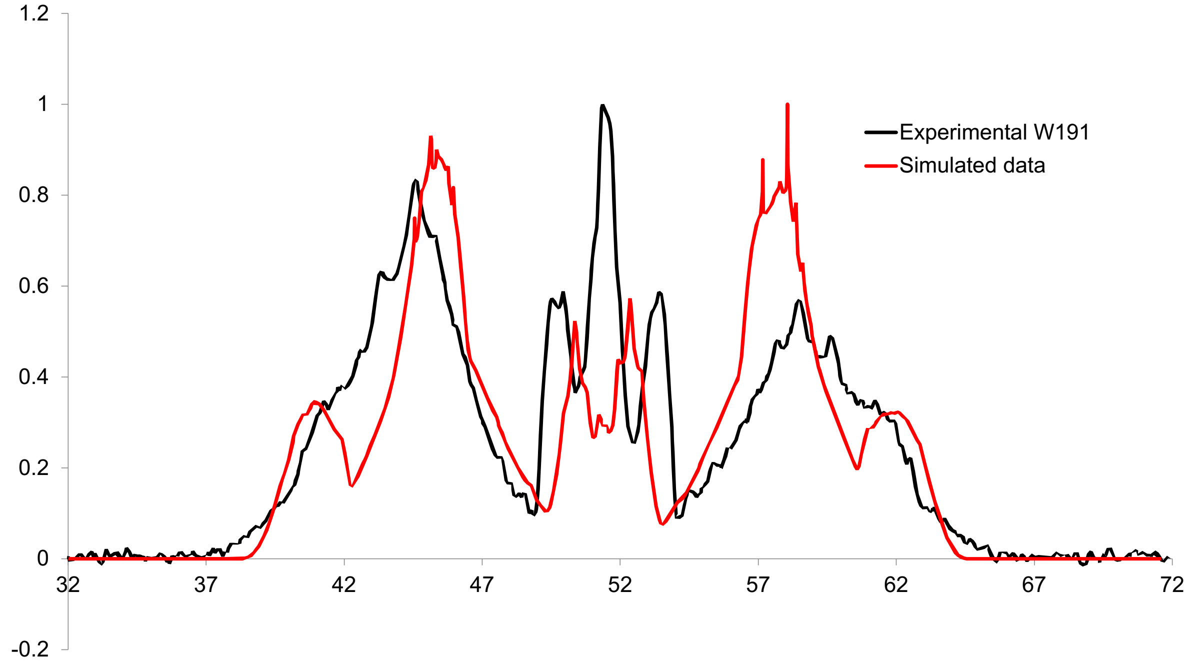


**Figure S12.** Experimental [1]H Davies ENDOR spectra (black) at 110 K for W191 ZnCcP with $CoN_5$ compared with the simulated spectra (red). The simulation was carried out using the saffron.m function of the easyspin matlab toolbox and the following g-tensors: $g_{xx}$ = 2.0033, $g_{yy}$ = 2.0024, $g_{zz}$ = 2.0023, and hyperfine coupling tensor reported by Bernini et al. (2014) of 7 protons were used for the simulation [2]: $^1H_{\beta 1}$ = [18.2, 19.4 26.3]; $^1H\beta_1$ = [9.6, 11.0, 16.2]; $^1H_1$ = [1.2, -14.4, -19.1]; $^1H_2$ = [1.7, -15.4, -20.2]; $^1H_5$ = [-6.7, -18.4, -24.7]; $^1H_6$ = [1.1, -3.4, -4.3]; $^1H_7$ = [ -4.6, -15.2, -22.4].

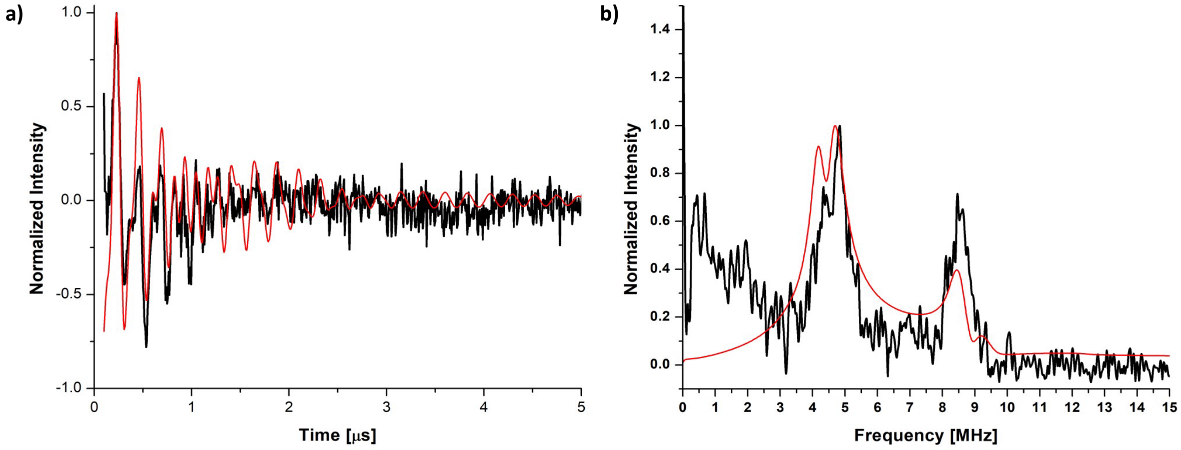


**Figure S13.** 3P-ESEEM experimental spectrum at 60 K of W191 ZnCcP (black) compared to simulated spectrum (red) **a)** in the time domain and **b)** in the frequency domain. The simulation was carried out with the following g-tensors: $g_{xx}$ = 2.0033, $g_{yy}$ = 2.0024, $g_{zz}$ = 2.0023 and the following hyperfine coupling constants of a single $^{14}N$ nucleus: $A_{xx}$ = -0.66 MHz, $A_{yy}$ = -0.87 MHz and $A_{zz}$ = +17.31 MHz. Spectra were simulated with the saffron.m function in EasySpin [3].

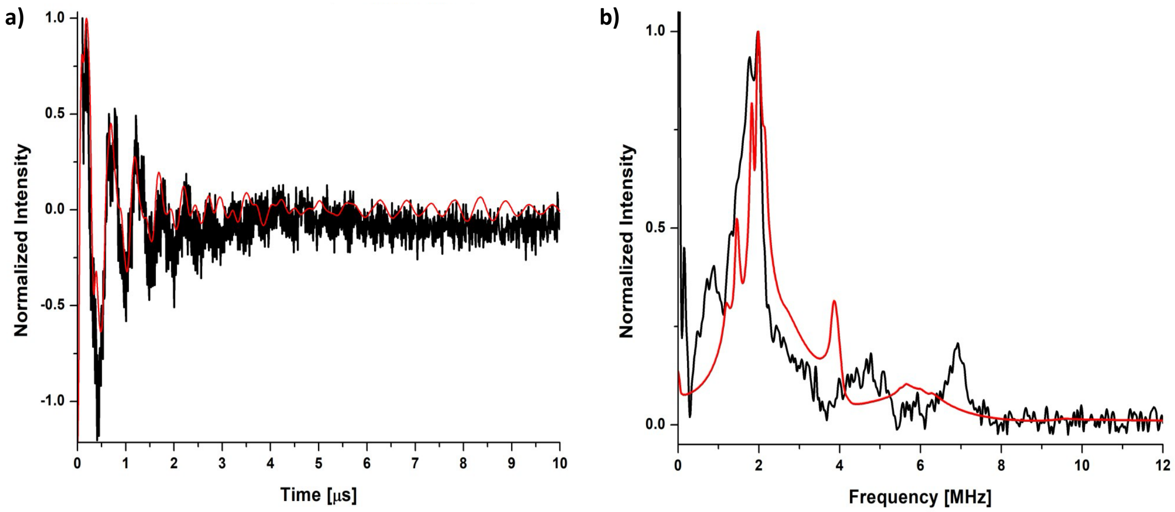


**Figure S14.** 3P-ESEEM experimental spectrum of Y191 ZnCcP (black) compared to simulated spectrum at 60 K (red) **a)** in the time domain and **b)** in the frequency domain. The simulation was carried out with the following g-tensors: $g_{xx}$ = 2.0033, $g_{yy}$ = 2.0024, $g_{zz}$ = 2.0023, the following hyperfine coupling constants of a single 14N nucleus: $A_{xx}$ = 3.677 MHz, $A_{yy}$ = 4.818 MHz and $A_{zz}$ = 11.709 MHz, and the following quadrupole coupling: $Q_{xx}$ = 0.085 MHz, $Q_{yy}$ = 0.216 MHz, $Q_{zz}$ = 0.096 MHz.

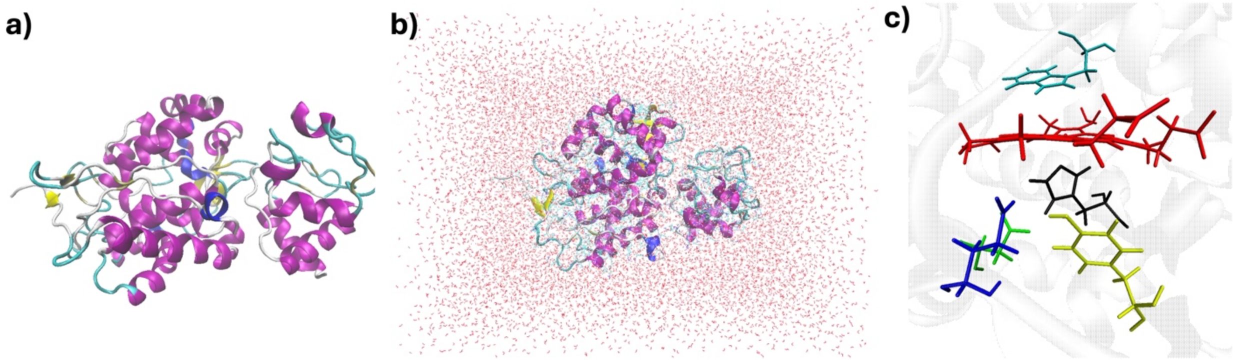


**Figure S15.** QM/MM Simulation details: **a)** Ribbon representation of the Y191:E232 structure (PDB ID: 6p41**) b)** Solvated protein in a rectangular TIP3P water box **c)** The QM region chosen for the QM/MM calculation. The residues are color coded: red (ZnP), yellow (Y191), black (H175), blue (E232), green (D235), cyan (W51).

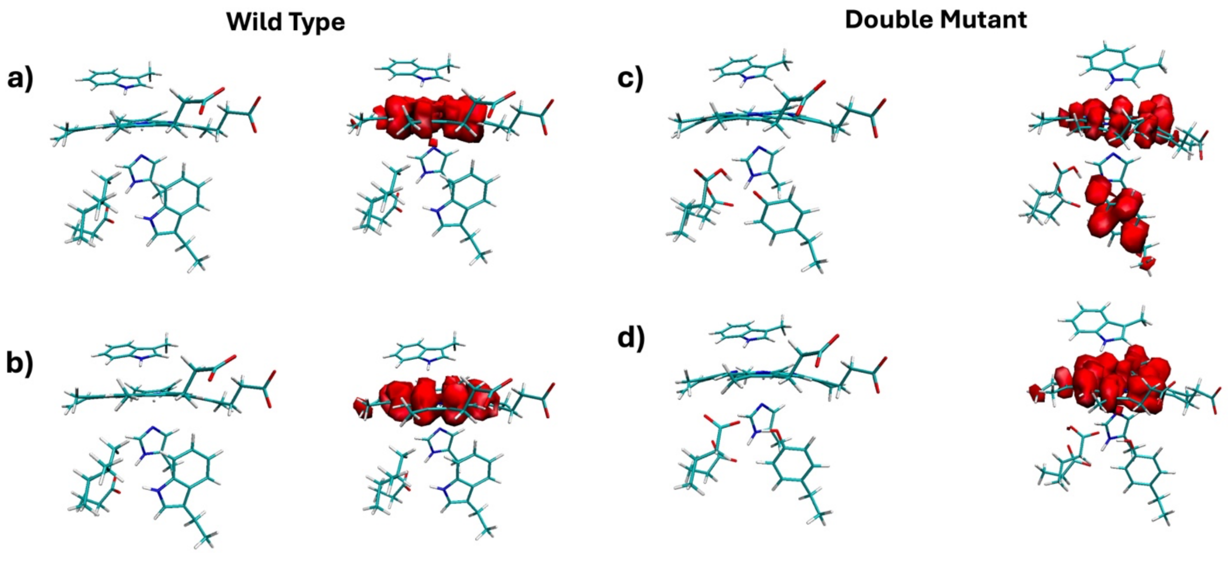


**Figure S16.** QM/MM optimized structures and spin density of Zn-triplet state: **a)** Optimized QM region of wild type CcP W191 with B3LYP/6-31g*. The distance between the H of the W191 residue and the O of the D235 is 1.82 Å. The spin density resides on ZnP **b)** Optimized QM region of wild type with wB97X/6-31g*. The distance between the H of the W191 residue and the O of the D235 is 1.80 Å. The spin density resides on ZnP **c)** Optimized QM region of Y191:E232 (Double Mutant) with B3LYP/6-31g*. In the optimized structure, the proton of the Y191 residue is transferred to the E232. The spin density resides both on ZnP and Y191 **d)** Optimized QM region of the Double Mutant with B3LYP/6-31g*. In the optimized structure, Y191 is protonated. The spin density resides on ZnP.

**Choice Of Functional:**

QM/MM optimization of the $^3$ZnCcP:Cc triplet state was performed with two different functionals – B3LYP and wB97x (Figure S5). For the wild type (WT), the QM/MM optimized structure for with both B3LYP and wB97x are similar, however, the $^3$ZnP SOMO from B3LYP has a positive energy whereas wB97x yields a negative energy as expected for a bound state. Similarly, for the

Y191:E232, B3LYP results in a positive energy state for the SOMO. For Y191:E232, B3LYP again yields a positive energy for the SOMO and an optimized structure where the proton transfers from the Y191 to the E232 residue. In addition, B3LPY yields spin density on both ZnP and Y191, which is unphysical for the $^3$ZnP triplet state. wB97x yields a negative SOMO energy, localized spin density on the $^3$ZnP and a protonated Y191 that is within H-bonding distance of the E232 residue. These results motivated our use of the range-separated hybrid functional wB97x in all QM/MM calculations reported in this manuscript.

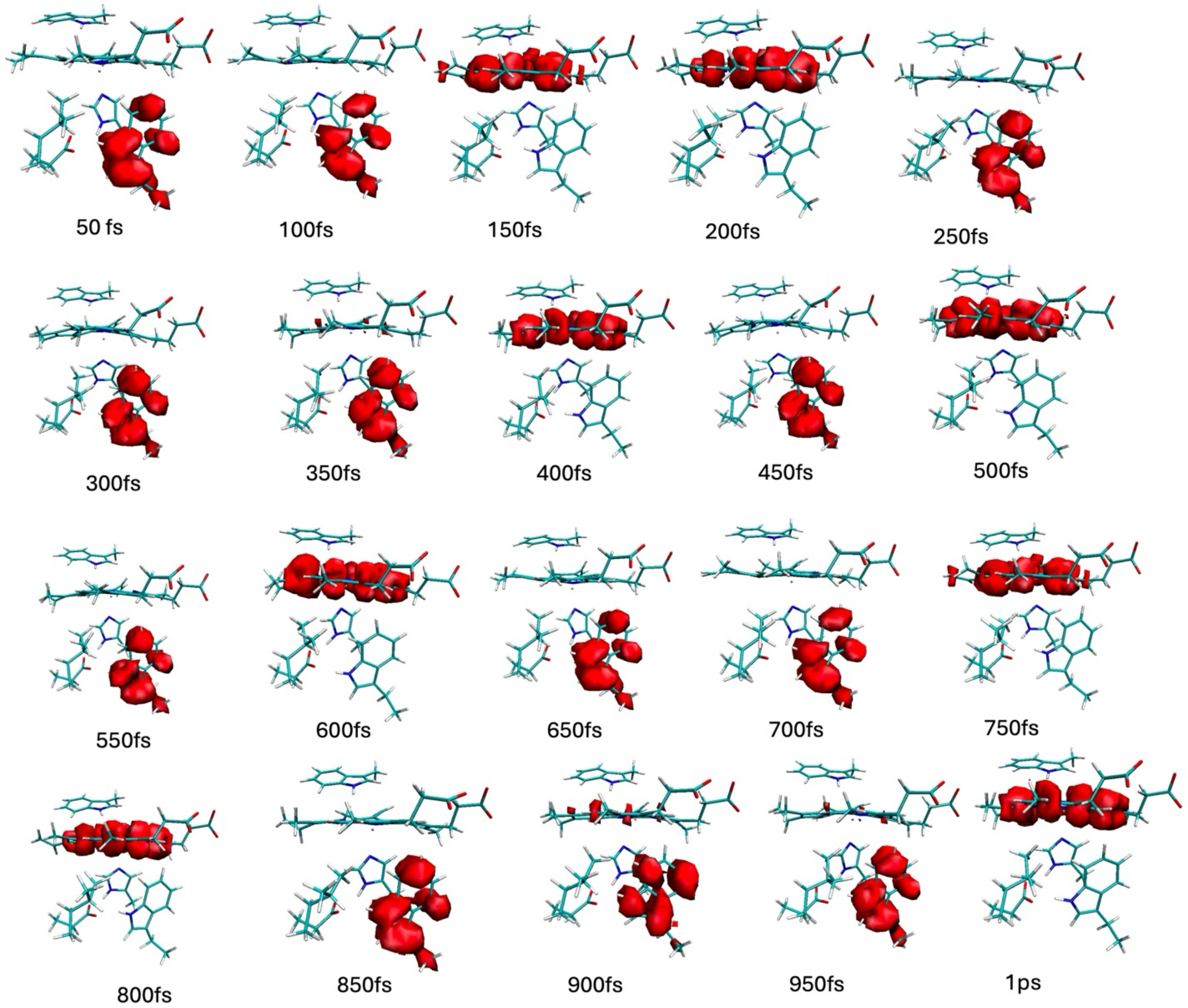


**Figure S17**. Spin density along the QM/MM NVT trajectory for the W191 calculated using wB97x/6-31g* for the QM region. The spin density along the trajectory shows alternating radical character between $ZnP^{\bullet+}$ and $Trp^{\bullet+}$. The hydrogen bond distance between the H of W191 and D235 remains within the range 1.6 Å -2.07 Å along the trajectory for the reported structures. The $K_{eq}$ lies toward $Trp^{\bullet+}$ unlike findings from previous computational studies [5], where the ESP charge fitting method is used on snapshots derived from MD trajectory for the radical location, which is approximate in case of electron density studies.

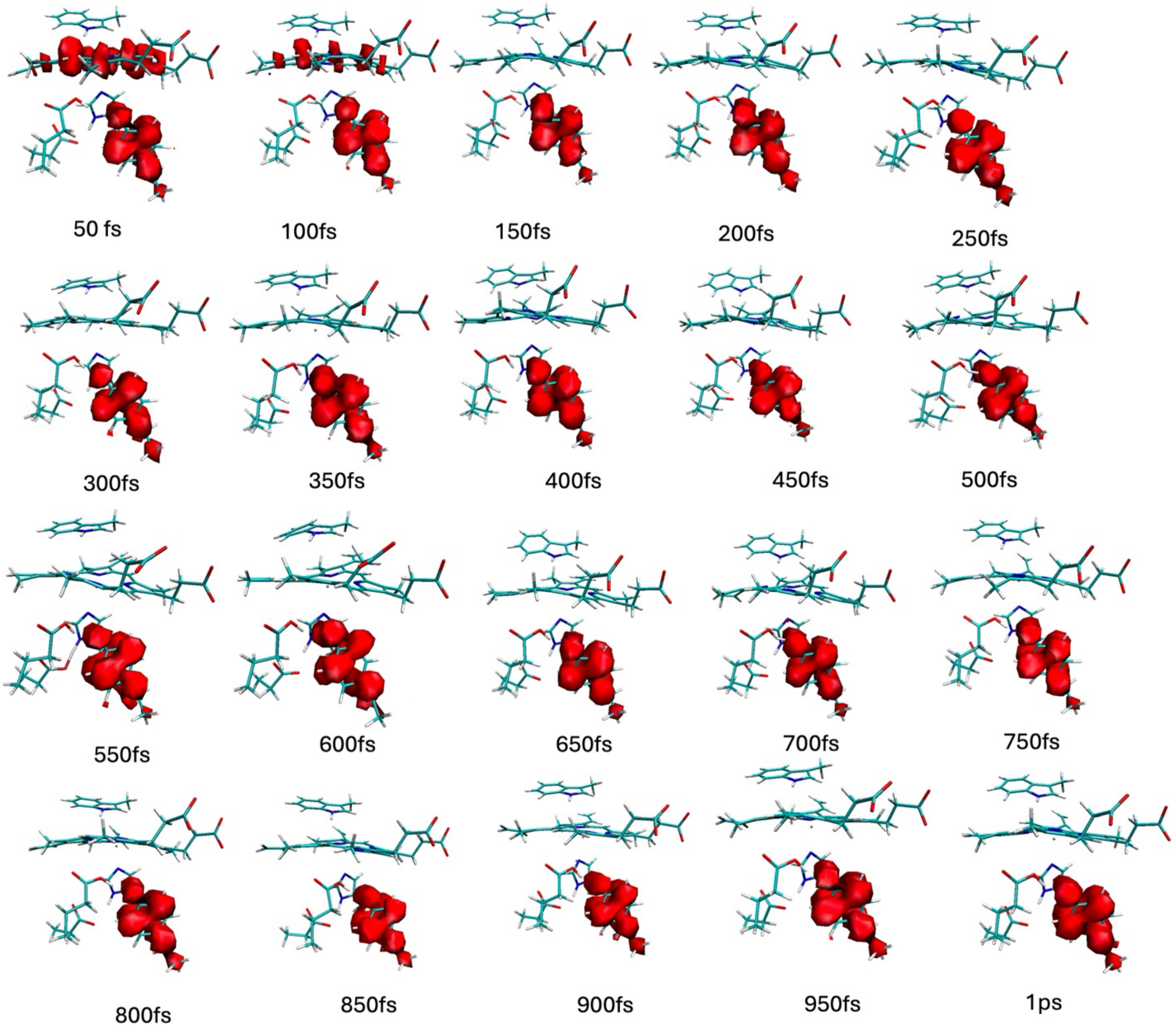


**Figure S18.** Spin density along the QM/MM NVT trajectory for the Y191:E232 calculated using wB97x/6-31g* for the QM region. The proton of Y191 is transferred to E232 in all of the structures. The spin density along the trajectory shows radical delocalized over ZnP and Y191 in the beginning and then the radical localizes to Y191 for the rest of the trajectory.

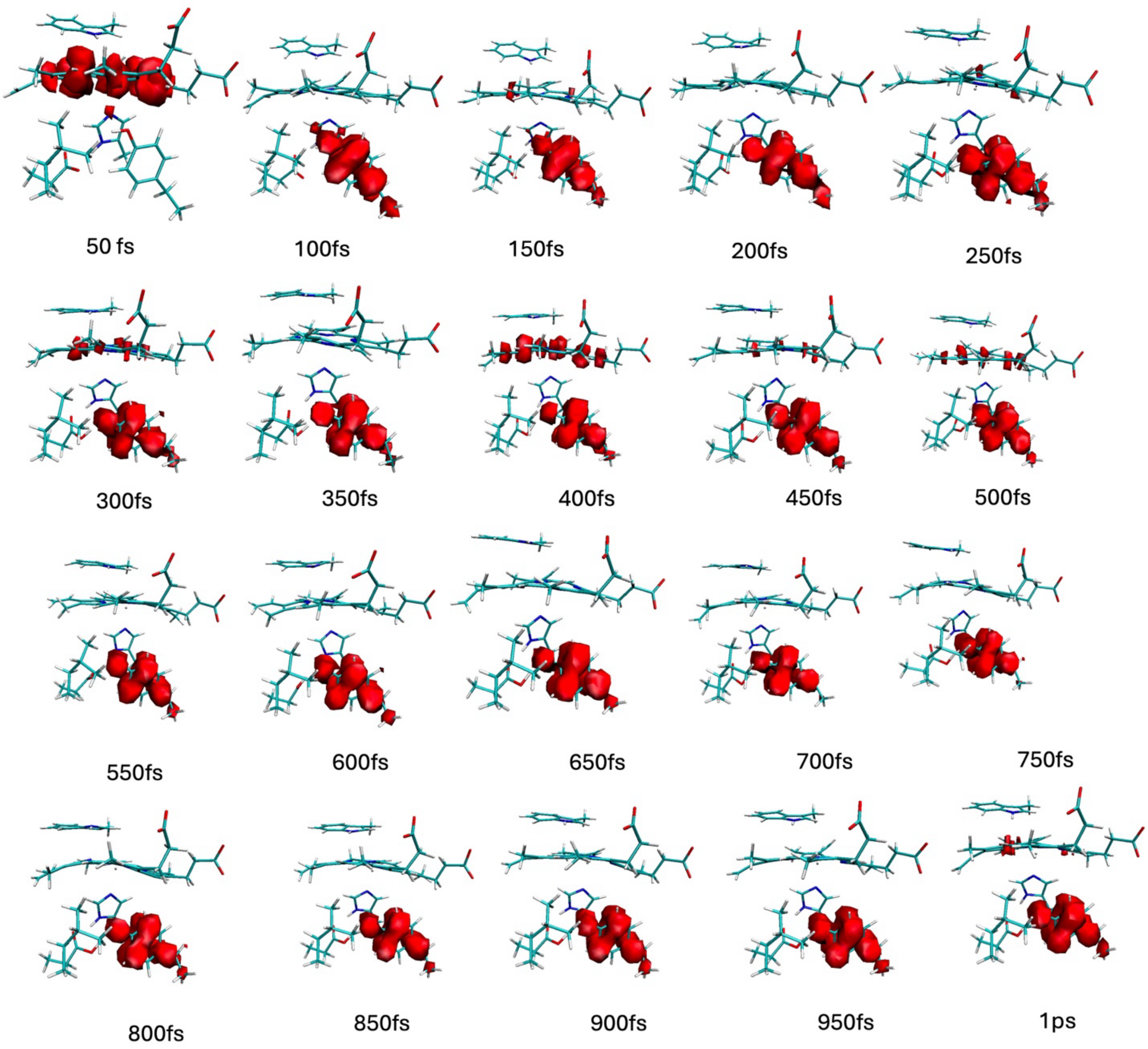


**Figure S19.** Spin density along the QM/MM NVT trajectory for Y191 calculated using wB97x/6-31g* for the QM region. The proton of the Y191 is transferred to the D235 from the structure at 300fs. The spin density along the trajectory shows radical mostly resides on Y191.

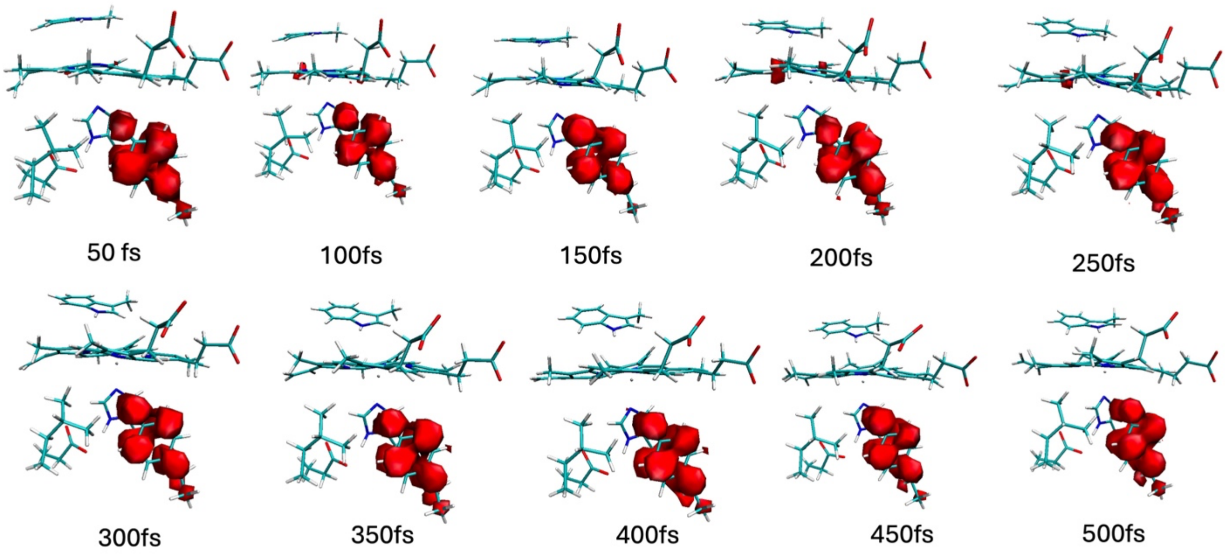


**Figure S20.** Spin density along the QM/MM NVT trajectory for Y191 with Y191 deprotonated calculated using wB97x/6-31g* for the QM region. The radical resides on Y191.

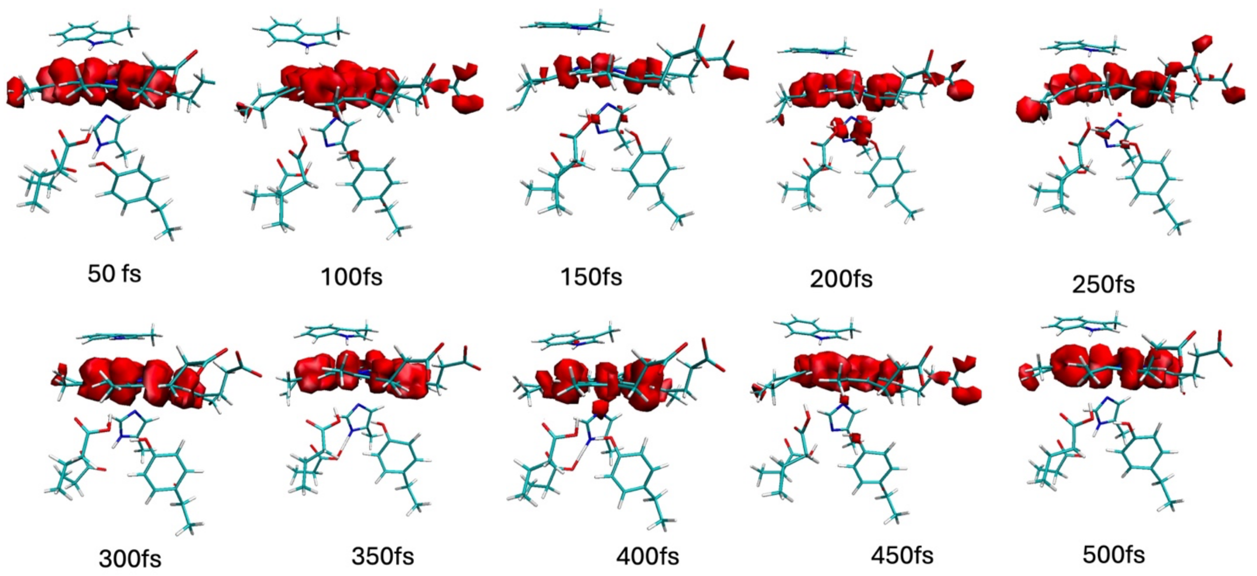


**Figure S21.** Spin density along the QM/MM NVT trajectory for Y191:E232 with E232 protonated calculated using wB97x/6-31g* for the QM region. The radical resides on ZnP.

**Github Folder:**

Github link: https://github.com/AnanthGroup/CcP-Ccstructurefile

- Pdb_files.zip: The PDB files 1u74, 6p41, 5cih used as the initial structure for the wild type, double mutant (Y191:E232), and single mutant (Y191) respectively.
- input_files.zip: This folder contains all the necessary input files for calculating the single point QM/MM calculation in CHEMSHELL for obtaining the spin density. The cm.psf file has all the charge and connectivity information for the protein. The top_all22_heme.rtf file contains the topology information and the file par_mod.prm contains the bonded and non-bonded parameters.
- WT_trajectory_structures.zip: This folder contains the structure coordinate of W191 in the chemshell input file format for all the snapshots along the QM/MM NVT 1ps trajectory, the optimized structure for triplet and doublet.
- DM_trajectory_structures.zip: This folder contains the structure coordinate of Y191:E232 in the chemshell input file format for all the snapshots along the QM/MM NVT 1ps trajectory, the optimized structure for triplet and doublet.
- SM_trajectory_structures.zip: This folder contains the structure coordinate of Y191in the chemshell input file format for all the snapshots along the QM/MM NVT 1ps trajectory, the optimized structure for doublet.
- DM_COOH.zip: This folder contains the structure coordinate of Y191:E232, E232 protonated in the chemshell input file format for the snapshots along the QM/MM NVT 500fs trajectory.

- SM_Tyr_deprotonated.zip: This folder contains the structure coordinate of Y191, Y191 protonated in the chemshell input file format for the snapshots along the QM/MM NVT 50fs trajectory
